# *Syzygium cumini* (L.) Skeels has Thermostable and Selective Anti-*Bacillus anthracis* Activity

**DOI:** 10.1101/2024.10.22.617365

**Authors:** Rajinder Kaur, Ashish Kumar Singh, Indresh Kumar Maurya, Samer Singh

**Affiliations:** Centre of Experimental Medicine and Surgery, Institute of Medical Sciences, Banaras Hindu University, Varanasi, 221005, India; Department of Microbial Biotechnology, Panjab University, Chandigarh, 160014, India

**Keywords:** Anthrax, Traditional Medicine, Medicinal plants, Aqueous Extract, Reemerging Disease

## Abstract

**Introduction:** Gastrointestinal (GI) anthrax caused by *Bacillus anthracis* remains a neglected disease in many parts of the Americas, Asia, and Africa. The symptoms include dysentery, stomachache, bloating of the stomach, vomiting, fever, chills, *etc.* Current study evaluated several edible plants traditionally indicated for GI disease/symptoms in the Indian subcontinent for their anti*-B. anthracis* activity.

**Materials & Methods:** Aqueous extracts of plant parts were assessed for anti-*B. anthracis* activity using standard antimicrobial susceptibility testing assays. Most promising extracts were evaluated for desirable activity under conditions relevant to their usage, including extremes of temperature, pH, presence of bile salt, impact on gut microflora, and interaction with FDA-approved drugs for anthrax treatment. The bioactive components separated by bioactivity-guided thin-layer-chromatography were subjected to GC-MS characterization.

**Results:** Aqueous *Syzygium cumini* (L.) Skeels or ‘Jamun’ extract (AJE) was most potent and reduced the viable colony-forming units (CFU) by 6-logs within 2 hours of exposure at ≥1.9%w/v concentration. It displayed both desirable selectivity towards gut microflora and thermostability (>90% and ∼80% of anti-*B. anthracis* activity were retained on incubation at 50°C for 20 days and at 95°C for 12h, respectively). AJE and FDA-approved antibiotics for anthrax displayed synergy. GC-MS analysis of AJE identified various previously-identified antimicrobials belonging to categories of alkaloids, flavonoids, phenols, *etc*.

**Conclusion:** AJE has potent and selective anti-*B. anthracis* activity with the desired degree of thermotolerance, compatibility with gut microflora, and recommended antibiotics. Further studies exploring the bioactive components in AJE and their potential application in preventing anthrax and anthrax-like diseases may be undertaken.

## 1. INTRODUCTION

Plants have always played a key role in combating ailments in livestock and humans in many indigenous communities (Bannerman 1980, 1982; Bussmann et al., 2011; WHO 2013). Globally, gastrointestinal infections/diseases remain a significant contributor to morbidity and mortality. Gastrointestinal (GI) anthrax in livestock and humans caused by the ingestion of *B. anthracis* spores from contaminated food or soil (Beatty et al., 2003; CDC, 2016, 2020) and generally characterized by dysentery (diarrhea or bloody diarrhea), stomachache, bloating of the stomach, vomiting, fever and chills, anorexia, nausea, abdominal pain, hematemesis and melena (Beatty et al., 2003; Owen et al., 2015), remains a major threat in endemic areas of Central and South America, southern Europe, Asia and Africa, parts of Australia (Shadomy et al., 2016; Turnbull, 2008; Carlson et al., 2019). The annual global incidence of human anthrax has been variously estimated at between 2000 and 20,000 cases per year (Amiri et al., 2021; Simonsen et al., 2022). It is occasionally reported from parts of Europe like France, Italy, the United Kingdom, *etc*. As late as 2019, anthrax topped the list of zoonotic diseases affecting the United Kingdom (Gov UK 2019). In the Indian subcontinent, anthrax is endemic. It is enzootic in southern India, but its incidence is less frequent or absent in northern India. In India, anthrax is endemic to the regions of Jammu and Kashmir, Orissa, Andhra Pradesh, Tamil Nadu, and Karnataka (ICAR-NIVEDI 2011). In the last six months, from January 2024 to June 2024, a total of 50 anthrax alerts were recorded from different parts of the world, such as India, Iraq, Germany, Kenya, Russia, Uganda, Armenia, Kazakhstan, Kyrgyzstan, China, and Ukraine (HealthMap 2024). Anthrax outbreaks in India over the past 15 years have affected at least 1,208 people and resulted in 436 deaths (Bhattacharya et al., 2019). During 2022–2023, eight outbreaks of anthrax in livestock were reported from different parts of India with a high rate of fatality (DAHD 2023).

The disease can clinically present itself as either an intestinal or oropharyngeal infection. Non-treatment or delays in treatment can be fatal (CDC, 2020). Traditionally, herbal extracts from different plant parts have been reported in both ethnoveterinary and ethnomedicinal practices for the treatment of various gastrointestinal (GI) conditions as indicated above, including anthrax in both animals and humans (Dold and Cocks, 2002; Merwe et al., 2001; Moshi et al., 2009). The use of plants such as *Senna italica* Mill., *Azadirachta indica* A. Juss.*, Teucrium africanum* Thunb., *Ptaeroxylon obliquum* (Thunb.) Radlk., *Achyrospermum schimperi* (Hochst. ex Briq.) Perkins, *Teucrium polium* L.*, Curcuma longa* L., and *Allium cepa* L. have been previously documented for anthrax (Darabpour et al., 2010; Dold and Cocks, 2002; Duke, 2008, 2002; Merwe et al., 2001; USDA, 2016). Recently, we have also provided evidence for the traditional use of *Allium sativum* L., *Allium cepa* L., *Azadirachta indica* A. Juss., *Berberis asiatica* Roxb. Ex DC., *Psidium guajava* L. and *Mangifera indica* L. against anthrax (Kaur et al., 2021).

The current study was performed to evaluate the purported anti-*B. anthracis* activity of nine commonly available livestock edible plants indicated in the traditional (folk and Ayurveda) medicine for dysentery (diarrhea or bloody diarrhea), stomachache, bloating of the stomach, vomiting, fever, chills, etc., symptoms commonly observed in gastrointestinal anthrax (see Table 1), to generate evidence for their traditional use. The identification of anti-*B. anthracis* activity of commonly available livestock edible plants was performed using *B. anthracis* Sterne 34F2 strain (*i.e*., pXO1^+^, pXO2^-^) that is used as veterinary vaccine, as surrogate for wild type (Bower et al., 2019; Kaur *et al*., 2013; Manish et al., 2020). We have tested the activity of plant parts of *Brassica nigra* (L) K. Koch. (Black mustard), *Zea mays* L. (Corn), *Triticum aestivum* L. (Wheat), *Syzygium cumini* L. Skeels (black plum or “Jamun”), *Polyalthia longifolia* (Sonn.) Thwaites (False Ashoka), *Ficus religiosa* L. (Sacred fig tree or “Peepal”), *Ficus benghalensis* L. (Banyan or “Bargad”), *Murraya koenigii* (L.) Spreng (curry leaf tree or “Sweet Neem”) and *Madhuca longifolia* (J.Koenig ex L.) J.F.Macbr (“Mahua”) against *B. anthracis.* These livestock edible plants are used as fodder and also employed to treat diarrhea, stomachache, and various GI microbial infections in livestock (Ayush 2015; Bussmann et al., 2011; Council of Scientific & Industrial Research (India), 1948; Dattatray et al., 2021; Duke, 2002, 2008, 1983; Millat et al., 2019; Hawaz Weldu et al., 2019;Yao et al., 2019) (Table 1).

**Table 1:**
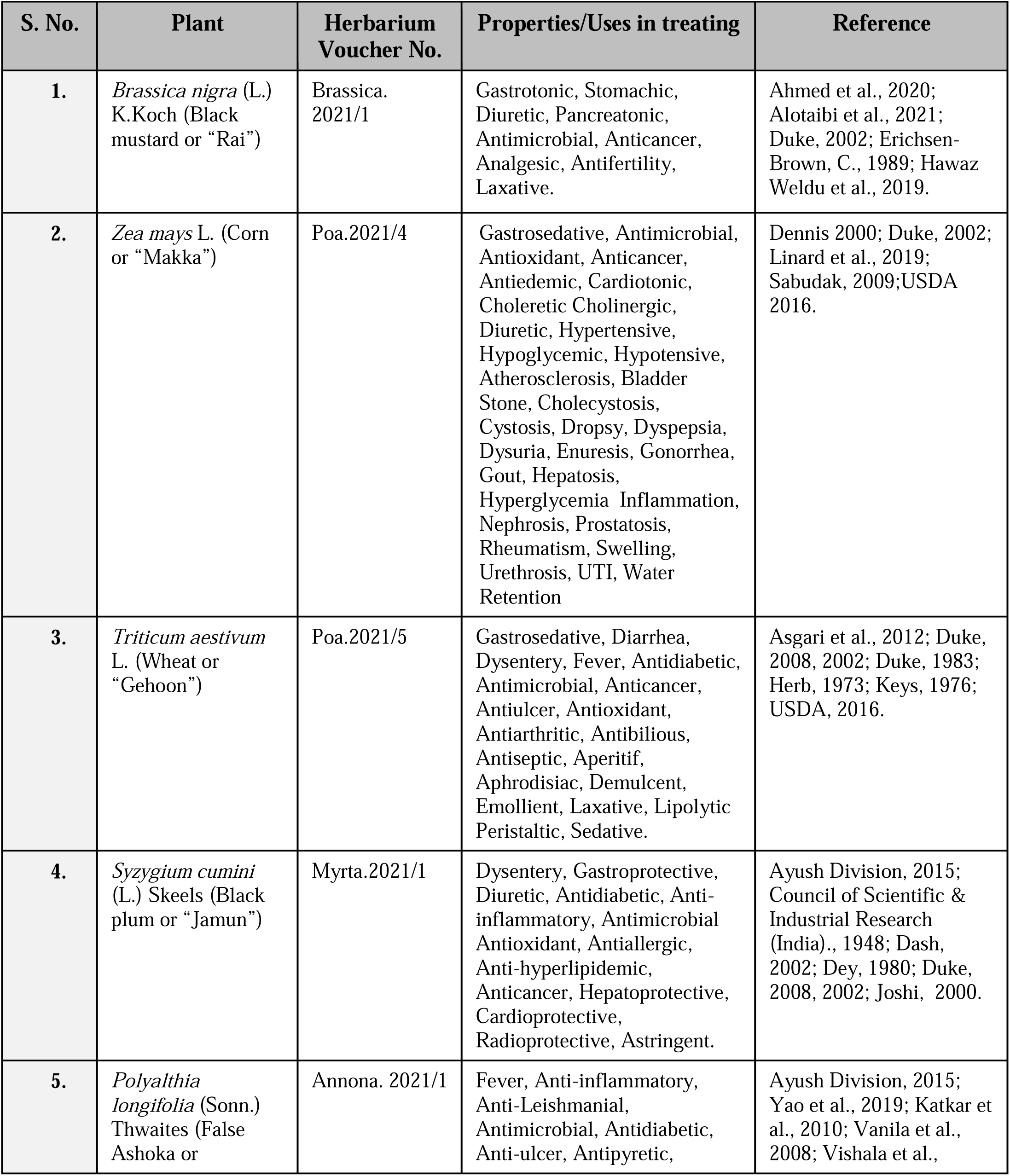
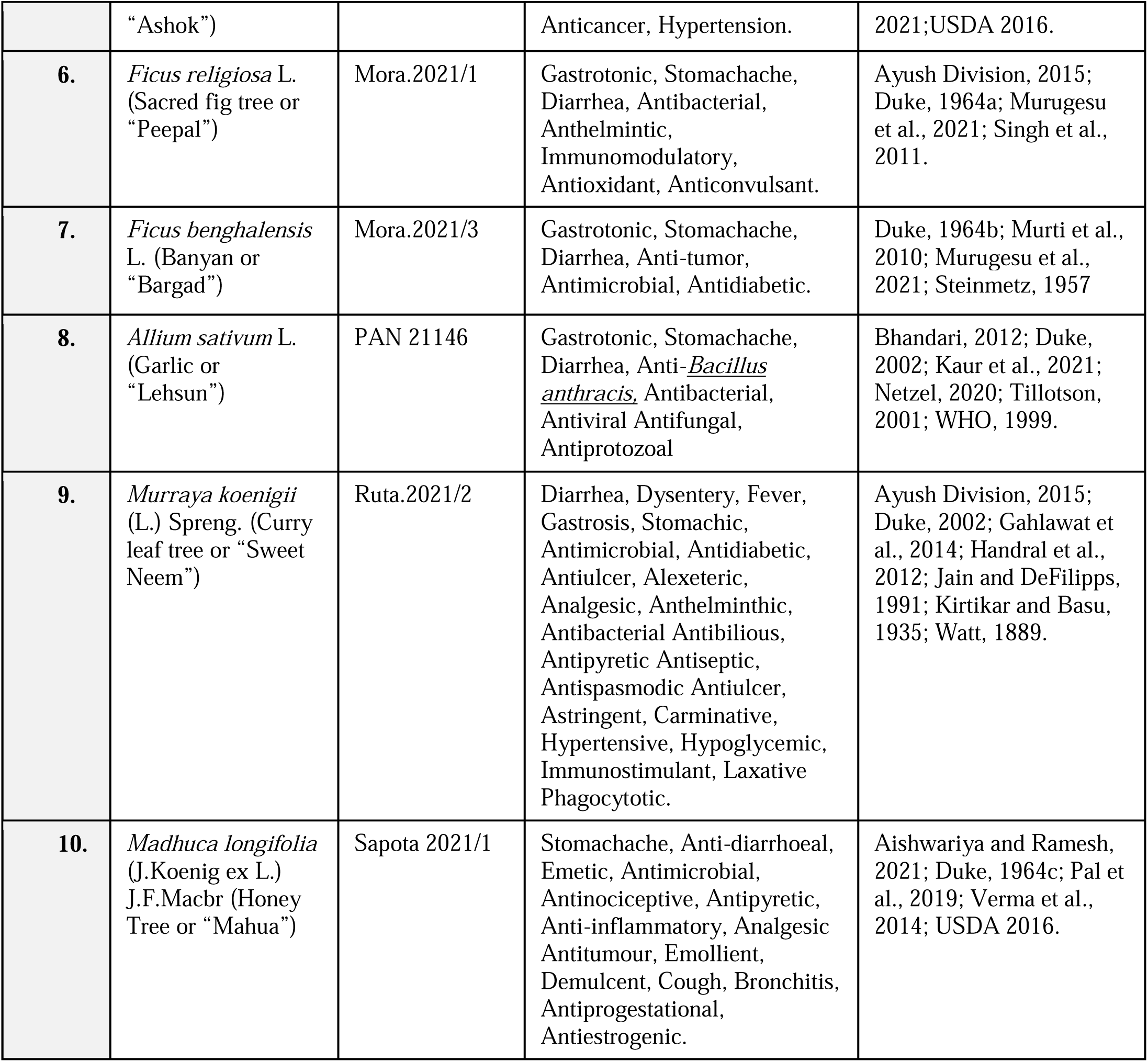
Properties and uses of plants used in the current study as indicated in literature.

Among the plants tested in the current study, *Syzygium cumini* L. Skeels (Jamun) displayed the most potent anti-*B. anthracis* activity. The exposure of *B. anthracis* to aqueous extract of *Syzygium cumini* L. Skeels leaves or aqueous Jamun extract (AJE) decreased the viable colony-forming unit (CFU) in broth by 6-logs within 2 h at a concentration of ≥1.9%w/v. Furthermore, it was not found to antagonize the activity of FDA-approved antibiotics for anthrax treatment suggesting its safer use in combination with antibiotics. The AJE showed more thermostability (heat stable) than the previously reported aqueous extract of *Allium sativum* L. (Garlic) (Kaur et al., 2021) and displayed a safer interaction with *Escherichia coli* (*E. coli*) a prominent commensal microflora of gastrointestinal tract in the presence of bile salt as well as other probiotic microflora. In Ayurveda medicine system the extracts from *Syzygium cumini* L. Skeels (Jamun) fruits, leaves, seeds, and bark have been reported for various GI diseases and indications (Ansari et al., 2021; Dash, 2002; Dey, 1980; Joshi and Joshi., 2000; Khalique and Rauf, 2016). Additionally, different parts of the Jamun are frequently indicated as anti-diarrheal, gastroprotective, anti-constipation, diuretic, anti-ulcerogenic, fever, gastropathy, stomachic, stomachalgia, antioxidant, anti-inflammatory, neuropsychopharmacological, antimicrobial, antibacterial, anti-HIV, antileishmanial, and antifungal, antifertility, anorexigenic, radioprotective, carminative, antiscorbutic, antidote for strychnine poisoning, skin diseases, leucorrhoea, etc. (Burkill, 1935; Duke, 2008; Jagetia, 2018; Quisumbing, 1951; Sagrawat, 2006; Swami et al., 2012; Dash, 2002; Dey, 1980; Joshi and Joshi., 2000; Khalique and Rauf, 2016). Local inhabitants of Rudraprayag district, part of western Himalaya, India use decoctions of Jamun bark for the treatment of dysentery and diarrhea (Singh et al., 2017).

## 2. MATERIAL AND METHODS

### 2.1. Bacterial strains and chemicals

In the current study, *B. anthracis* Sterne 34F2 – an avirulent (pXO1^+^, pXO2^-^) vaccine strain and *Escherichia coli* DH5α (*E. coli* DH5α) were used as surrogate for wild type *B. anthracis* and commensal *E. coli*, respectively. Mueller Hinton Broth (MHB), Bacteriological agar powder, Mueller Hinton Agar (MHA), Normal saline solution (NSS), and MTT (3-(4,5-dimethylthiazol-2-yl)-2,5-diphenyltetrazolium bromide) (used for cell viability assay) were purchased from HiMedia Laboratories Ltd. (India). The antibiotics and organic solvents used were purchased from Sigma Aldrich Inc (St. Louis, MO, USA) and Merck Millipore (Merck Life Science Private Limited, India), respectively.

### 2.2. Collection of plant samples

Fresh leaves from identified plants [*i.e., Brassica nigra* (L.) K. Koch (Black mustard), *Zea mays* L. (Corn), *Triticum aestivum* L. (Wheat), *Syzygium cumini* (L). Skeels (black plum or “Jamun”), *Polyalthia longifolia* (Sonn.) Thwaites (False Ashoka), *Ficus religiosa* L. (Sacred fig tree or “Peepal”), *Ficus benghalensis* L. (Banyan or “Bargad”), *Murraya koenigii* (L.) Spreng. (Curry leaf tree or “Sweet Neem”) and *Madhuca longifolia* (J.Koenig ex L.) J.F.Macbr. (“Mahua”)] were from the premises of Banaras Hindu University, Varanasi, India. Fresh leaves from all the selected plants were collected in sterile polythene bags on the day experiments were performed. The dried *Allium sativum* L. bulbs were purchased from sector 15, Chandigarh, India (Table 2).

**Table 2:** List of plants evaluated for anti-*B. anthracis* activity along with the average of measured Zone of inhibition.

| S. No. | Sample (Conc. w/v) | Part used | Collection location | Vol. loaded in wells | Zone of inhibition (mm) |  |  |
| --- | --- | --- | --- | --- | --- | --- | --- |
|  |  |  |  |  | 24hr | 48hr | 72hr |
| 1. | *Brassica nigra* (L.)<br>*K.Koch* (Black mustard or “Rai”) (80% w/v) | Leaf | BHU, Varanasi | 60µL | 0 | 0 | 0 |
| 2. | *Brassica nigra* (L.)<br>*K.Koch* (Black mustard or “Rai”) (80% w/v) | Flower | BHU, Varanasi | 60µL | 0 | 0 | 0 |
| 3. | *Zea mays* L. (Corn or “Makka”) (80% w/v) | Leaf | BHU, Varanasi | 60µL | 0 | 0 | 0 |
| 4. | *Triticum aestivum* L. (Wheat or “Gehoon”) (80% w/v) | Leaf | BHU, Varanasi | 60µL | 0 | 0 | 0 |
| 5. | *Syzygium cumini* L. Skeels (Black plum or “Jamun”) (80% w/v) | Leaf | BHU, Varanasi | 60µL | 16 | 15 | 14 |
| 6. | *Polyalthia longifolia* (Sonn.) Thwaites (False Ashoka or “Ashok”) (80% w/v)* | Leaf | BHU, Varanasi | 60µL | 13 | 13 | 12 |
| 7. | *Allium sativum* L. (Garlic or “Lehsun”) (40% w/v) | Cloves/Bu lb | Local market, Chandigarh | 60µL | 25 | 20 | 20 |
| 8. | *Cynodon dactylon* L. (Bermuda grass) (80% w/v) | Leafs | BHU, Varanasi | 60µL | 0 | 0 | 0 |
| 9. | *Ficus religiosa* L. (Sacred fig tree or “Peepal”) (80% w/v) | Leafs | BHU, Varanasi | 60µL | 0 | 0 | 0 |
| 10. | *Ficus benghalensis* L. (Banyan or “Bargad”) (80% w/v) | Leafs | BHU, Varanasi | 60µL | 0 | 0 | 0 |
| 11. | *Madhuca longifolia* (J.Koenig ex L.)<br>J.F.Macbr (Honey tree or “Mahua”) (80% w/v) | Leaf | BHU, Varanasi | 60µL | 12 | 10 | 0 |
| 12. | *Madhuca longifolia*<br>(J.Koenig ex L.)<br>J.F.Macbr (Honey tree<br>or “Mahua”) (80% w/v) | Fruit | BHU,<br>Varanasi | 60µL | 0 | 0 | 0 |
| 13. | *Murraya koenigii* (L.)<br>Spreng (curry leaf tree<br>or “Sweet Neem”) (80%<br>w/v) | Leafs | BHU,<br>Varanasi | 60µL | 0 | 0 | 0 |
| 14. | Positive ctrl.<br>(Rifampicin) | - | Himedia | 8µg | 20 | 20 | 20 |
| 15. | Negative ctrl. (Water) | - | Autoclaved<br>distilled water | 60µL | 0 | 0 | 0 |

\*used concentration did not show any activity in liquid /broth medium.

### 2.3. The aqueous plant extract preparation

The freshly collected leaves, fruits, and garlic cloves were washed with distilled water to remove any dust particles followed by surface disinfection using 70% ethanol (Kaur et al., 2021). Samples were dried at room temperature for up to 15 min to remove any ethanol traces from them. Plant samples (*i.e.,* leaves, fruits, and garlic clove) were weighted (*e.g.,* 2-4 g) and placed in sterile mortar pestles where they were crushed in sterile ultrapure water (*e.g.,* 5-10 mL). The aqueous soluble extract or soup was separated from the plant debris by passing the crushed samples through a sterile filter paper (*i.e.,* Whatman No. 1). The separated soup or aqueous plant extract was collected in a sterile 15 mL tube and stored at 4°C until its further use. The strength of aqueous plant extract stocks prepared was calculated as weight of the fresh plant material to volume of the extractant water used for the sample/stock preparation and reported as % w/v, (*e.g.*, 400 mg leaf used to prepare 1 mL of aqueous extract is reported as 40% w/v or 400 mg/ml (Kaur et al., 2021). The final concentration of aqueous plant extracts in the test cultures was achieved by its dilution in the suitable medium or the indicated diluent in experiment.

### 2.4. Assessing the anti-*B. anthracis* activity of aqueous plant extracts

The anti-*B. anthracis* activity of all the selected aqueous plant extracts was assessed by Agar Well Diffusion Assay (AWDA) as indicated in the CLSI guidelines (CLSI, 2016) using the *B. anthracis* Sterne 34F2 strain as a surrogate for wild type organism. The assay was performed essentially as reported in our previous publication (Kaur et al., 2021). Briefly, the broth dilution assay (Minimum inhibitory concentration (MIC) determination assay) was performed by inoculating 0.1 mL of an overnight grown culture of *B. anthracis* in 5 mL of fresh MHB medium and incubated at 37°C with shaking (120 rpm) for 2–3 h until it reached 0.4 optical density (OD) at 600 nm (OD_600nm_). The broth culture of 0.4 OD_600nm_ was further diluted in MHB medium to 0.1 OD_600nm_ (app. 1-2 × 10^6^ CFU/mL). In 96 well flat-bottom plates different volumes of 80%w/v AJE were added in wells containing 200µL of 0.1 OD_600nm_ *B. anthracis* broth cultures, so as to achieve the final concentration of AGE in the range of 0 to 20% w/v. The plates were incubated at 37°C with shaking at 120 rpm for 12-16 h for ascertaining its impact on their growth.

### 2.5. Evaluating bacteriostatic and bactericidal activity of aqueous *Syzygium cumini* L Skeels (Jamun) extract (AJE)

The bactericidal or bacteriostatic activity of the AJE was assessed by performing a time-kill assay (CLSI, 2016). The inoculated *B. anthracis* culture of approximately 0.4 OD_600nm_ was centrifuged at 8000g for 5 min to pellet down the *B. anthracis* cells. The cell pellets were washed with normal saline solution (0.85% NaCl) to remove any media traces left, followed by suspension in NSS or MHB media. Approximately 1 × 10^6^ CFU/mL (*i.e.,* 0.1 OD_600nm)_ of *B. anthracis* was supplemented with AJE at a final concentration of 0–3.6% w/v and incubated at 37°C with shaking (120 rpm) for 0-24 h. After every 2 h intervals, AJE exposed *B. anthracis* culture was serially diluted and plated on MHA medium plates, and the plates were incubated at 37°C for 15–18 h. The number of colonies obtained at different time intervals was counted and used to calculate the residual CFU/mL at different time durations (0–24 h) of exposure to different concentrations of AJE.

### 2.6. Morphological characterization of AJE exposed *B. anthracis* cells by Scanning Electron Microscopy (SEM)

The exponentially growing *B. anthracis* cells were exposed to a final concentration of 0-3.6% w/v of AJE for 2 h followed by processing for observation by SEM, essentially as described previously (Kaur et al., 2021). Briefly, the *B. anthracis* cells were fixed with 2% glutaraldehyde for 1 h at room temperature, washed with phosphate buffer (1X, pH 7.2), dehydrated by serial passage in 10, 30, 50, 75, 90, and 100% ethanol and finally sputter coated with gold. The images were taken at different magnifications using a ZEISS Scanning Electron Microscope (Department of Geology, Banaras Hindu University, Varanasi, India).

### 2.7. Activity modification of AJE by bile salt and comparison with *E. coli* and other commensal gut microbes present in commercially available probiotics

The comparative antibacterial activity of AJE to inhibit *B. anthracis, E. coli* DH5α, and gut microbes present in two different commercial probiotics formulations [*i.e.*, *Streptococcus faecalis, B. mesentericus, Clostridium butyricum, Lactobacillus sporogenes, Lactobacillus acidophilus, Lactobacillus plantarum, Lactobacillus casei, Lactobacillus rhamnosus, Streptococcus thermophiles, Saccharomyces boulardii, Bifidobacterium longum, Bifidobacterium breve, and Bifidobacterium infanti*] was tested with and without sodium deoxycholate supplementation (0.0025%) by AWDA essentially as described earlier (Kaur et al., 2021). Briefly, to test the relative potency of AJE against test microbes, 80μl of 80% w/v AJE stock was loaded in the well punched on the midline of two halves of the MHA plate (spread with *B. anthracis* and *E. coli* DH5α or the test probiotics formulations, in halves). The rifampicin (8μg/well) and ultrapure water were used as positive and negative/ solvent control, respectively. The plates were incubated at 37°C for 15-18 h. The zone of inhibition (ZOI) was measured and compared to assess the relative antibacterial activity of AJE against test organisms.

### 2.8. Effect of the extreme pH conditions, encountered in the gastrointestinal tract, on anti-*B. anthracis* activity of AJE

Effect of conditions encountered in the gastrointestinal tract, *i.e.,* extremes of pH on the anti-*B. anthracis* activity of AJE was assessed by AWDA (Kaur et al., 2021). Briefly, the pH of AJE (80% w/v) aliquots was adjusted in the range of 2–8 using 1N HCl or 1N NaOH and incubated at 37°C for different time durations (0–4 h). The 100μl aliquots of the incubated AJE samples were assayed for their residual anti-*B. anthracis* activity. The controls included AJE without any pH adjustment and normal saline solutions of matching pH (2, 3, and 8) as negative controls (– Ctrl). The ZOI produced by different samples was measured after overnight incubation of MHA plates at 37°C.

### 2.9. Interaction of FDA-approved antibiotics for anthrax treatment with AJE

The effect of FDA-approved antibiotics recommended for anthrax treatment, *i.e.,* Amoxicillin, Ciprofloxacin, Doxycycline, Levofloxacin, Penicillin, Rifampicin, and Tetracycline (CDC, 2016; U.S. Food and Drug Administration, 2018), on the anti-*B. anthracis* activity of AJE when used in combination was determined by minimum inhibitory concentration (MIC) determination assay, and the fractional inhibitory concentration index (FICI) for combinations was estimated as reported (Kaur et al., 2021).

### 2.10. Interaction of AJE with virulence plasmid pXO1 of *B. anthracis* (Virulence plasmid loss assay)

The potential of AJE to cause virulent plasmid (pXO1) loss from *B. anthracis* cell, after its treatment with AJE at its sub-lethal concentrations (*i.e.,* ≤ 1% w/v) was assessed. The plasmid curing was checked by performing colony PCR of the colonies obtained on MHA plates by plating *B. anthracis* cells exposed to AJE for different durations. The PCR conditions and the primers used were as previously described (Kaur et al., 2021).

### 2.11. Thin Layer Chromatography (TLC) followed by Bioautography and GC-MS analysis of AJE

TLC was performed in duplicates on silica gel 60 (Merck Millipore). The AJE extract was spotted 1 cm away from the base of the plate and run using the Toluene and Acetone (7:3 ratio) solvent system unless noted otherwise. The plates were dried and visualized under UV light. From one chromatogram the bioactive band obtained were scraped from the plate and dissolved in acetone. The mixture was then centrifuged to separate silica and the supernatant was used for GC-MS analysis whereas the other chromatogram was used for bioautography. The detailed procedure adopted was as published (Kaur et al., 2021).

### 2.12. Thermal stability of AJE

The thermostability of AJE and AGE anti-*B. anthracis* activity at high temperatures was determined for comparison. The 200μl aliquots of AGE (40% w/v) and AJE (80% w/v) were kept at 4°C-50°C for 0 h to 20 days. After incubation, the residual activity of the extract aliquots was evaluated by AWDA. The thermal stability at a still higher temperature was evaluated by incubating the 200 μl aliquots of AGE (40% w/v) and AJE (80% w/v) at 95°C for 0 h to 12 h, followed by residual activity determination by AWDA. The observed ZOI was measured, and the relative ZOI (%) was calculated [the relative ZOI (%) = Observed ZOI for sample incubated at test temperature (°C)/Observed ZOI for fresh aqueous extract *100]. (Note: As ZOI produced by 40% w/v of AJE aliquots was smaller than that of 40% w/v of AGE, a higher concentration stock of AJE (i.e., 80% w/v) was employed for activity comparison in AWDA.)

### 2.13. Data collection, analysis and statistics

Each experiment was performed in triplicates and repeated at least three times unless noted otherwise. The values shown are the average of independent experiments. Error bars represent the standard error of the mean (S.E.M.).

## 3. RESULTS

### 3.1. AJE has potent anti-*B. anthracis* activity

The anti-*B. anthracis* activity of tested aqueous plant extracts from livestock edible plants is summarized in Table 2. The aqueous extracts of *Syzygium cumini* L. Skeels and *Polyalthia longifolia* (Sonn.) Thwaites inhibited the growth of *B. anthracis* in AWDA (Fig. 1a). Among the tested extracts, aqueous *Syzygium cumini* L. Skeels extract (AJE) has displayed the highest and most potent anti-*B. anthracis* activity. The 16 mm diameter zone of inhibition (ZOI) produced by AJE was sustained for 72 h of incubation at 37°C similar to that produced by Rifampicin (+ Ctrl), indicative of bactericidal activity. Although, the aqueous leaf extracts of *Brassica nigra* (Mustard), *Zea mays* (Corn), *Triticum aestivum* (Wheat), *Cynodon dactylon* (Bermuda grass), *Ficus religiosa* (Sacred Fig tree or ‘Peepal’), *Ficus benghalensis* (Banyan), *Madhuca longifolia* (Mahua) and *Murraya koenigii* (Sweet Neem), have been indicated for treating various microbial infections, they did not show anti-*B. anthracis* activity at the tested concentration in both liquid and solid growth medium assays (Fig. 1b-c). The head-to-head comparison of the anti-*B. anthracis* activity of different aqueous extracts of the current study with that of *Allium sativum* L. (AGE), previously shown by us (Kaur et al., 2021), indicated the potency of anti-*B. anthracis* activity to be present in the decreasing order on a w/v basis as, *Allium sativum* L. (AGE)> *Syzygium cumini* L. Skeels (AJE)> *Polyalthia longifolia* (Sonn.) Thwaites (AAE) (Fig. 1a-c). Among the tested plant extracts in the current study, *Syzygium cumini* extract showed maximum potency and minimum batch-to-batch variations between experiments on a weight-to-weight basis, so it was selected for further characterization of its anti-*B. anthracis* activity. It is pertinent to mention here that different solvent extracts of *Syzygium cumini* L. Skeels leaf (*i.e.,* methanol, chloroform, acetone, hexane, butanol, and ethyl acetate) were also evaluated for their anti-*B anthracis* activity (Suppl. Fig. 1a). Methanol and acetone extracts displayed slightly higher activity as compared to the aqueous extract, as per the ZOIs produced. Interestingly, the dried residues of methanol and acetone extracts on reconstitution in water displayed *B. anthracis* activity (Suppl. Fig. 1b) suggesting the polar nature of compounds that displayed anti-*B. anthracis* activity in these extracts. As we preferred water (aqua) as an extractant for its straight forward field applicability, unlike other solvents, further characterization of the aqueous extract was performed.

**Figure 1.**
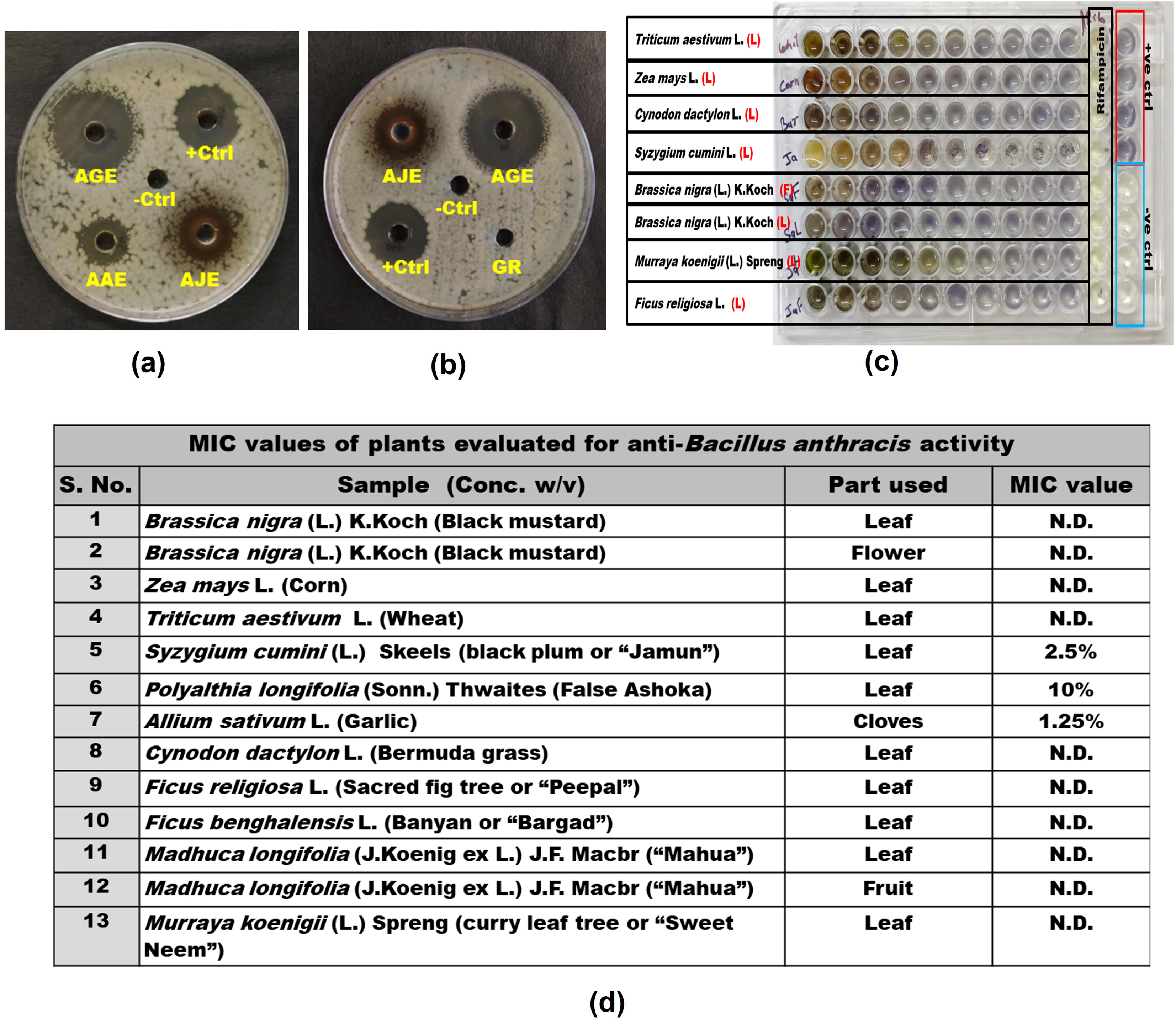
Common edible plants have anti-*B. anthracis* activity. (**a-i-ii**) The ZOI produced by different aqueous plant extracts in agar-well diffusion assay (AWDA) after 24 h incubation at 37°C [AJE: *Syzygium cumini L* (Black plum or “Jamun”) Leaf; AGE: *Allium sativum* L. (Garlic) clove; GR: *Cynodon dactylon* L. (common Bermuda grass); AAE: *Polyalthia longifolia S.* (False Ashoka) Leaf; +Ctrl: Antibiotic Rifampicin; –Ctrl: ultrapure water]. After AGE, AJE displayed the most potent anti-*B. anthracis* activity followed by AAE (See Table 2); **(b)** Comparative ZOI of AJE, AGE, GR when incubated at 37°C; **(c)** Minimum inhibitory concentration (MIC) determination assay: The supplementation of exponentially growing culture (1 x 10^6^ CFU/mL) cells with different concentration of aqueous extracts inhibited *B. anthracis* cells in a dose-dependent manner; **(d)** The MIC values shown are the average of three independent experiments (n = 3). N.D. – Not detected at the tested concentrations *i.e.,* 20% w/v.

### 3.2. AJE inhibits the growth of *B. anthracis* in a dose-dependent manner

The supplementation of exponentially growing *B. anthracis* cells in MHB growth medium (*i.e.,* 2 x 10^6^ CFU/mL) with different concentrations of AJE, *i.e.,* 0-20% w/v, was found to inhibit the growth of *B. anthracis* cells in a dose-dependent manner. It was observed that the minimum inhibitory concentration (MIC) of AJE for *B. anthracis* cells (when kept without any agitation or shaking in a 96-well flat bottom plate) was ≥2.5% w/v, whereas for *Polyalthia longifolia* it was 10% w/v (Fig. 1c-d). Extracts from other plants did not inhibit the growth of *B. anthracis* in the tested concentration range.

### 3.3. AJE is bactericidal to vegetative *B. anthracis* cells

The bactericidal activity of AJE was assessed by exposing exponentially growing *B. anthracis* cells with different concentrations of AJE (*i.e.,* 0-3.6% w/v) in the presence and absence of growth medium, *i.e.,* MHB and NSS, respectively, followed by estimating the number of residual colonies forming units (CFU)/mL at different time intervals (0–24 h). In the presence of enriched media, *i.e.,* MHB, the viability of vegetative *B. anthracis* cells in the presence of AJE at ≥1.9% w/v final concentration was found to decrease by 6 logs (Fig. 2a) within 2 h of incubation (agitation at 120 rpm) at 37°C. However, at lower concentrations (i.e., 1% w/v AJE), the bactericidal action on *B. anthracis* was delayed, and it took about 4 hours to kill all vegetative cells in the culture. In the absence of nutrient rich media, *i.e,* normal saline or NSS, the exposure to AJE at a final concentration of ≥1% w/v was able to kill *B. anthracis* cells completely within 2 h of incubation (Fig. 2b). Together, it suggests increased efficacy of AJE in killing vegetative *B. anthracis* cells in the absence of an organic medium. [*Note*: Based on the dry weight of residues present in the aqueous extract, the calculated concentrations of AJE residues present in 1% w/v, 1.9% w/v, and 3.6% w/v would be 1.12 mg/mL, 2.13 mg/mL, and 4.04 mg/mL, respectively.]

**Figure 2.**
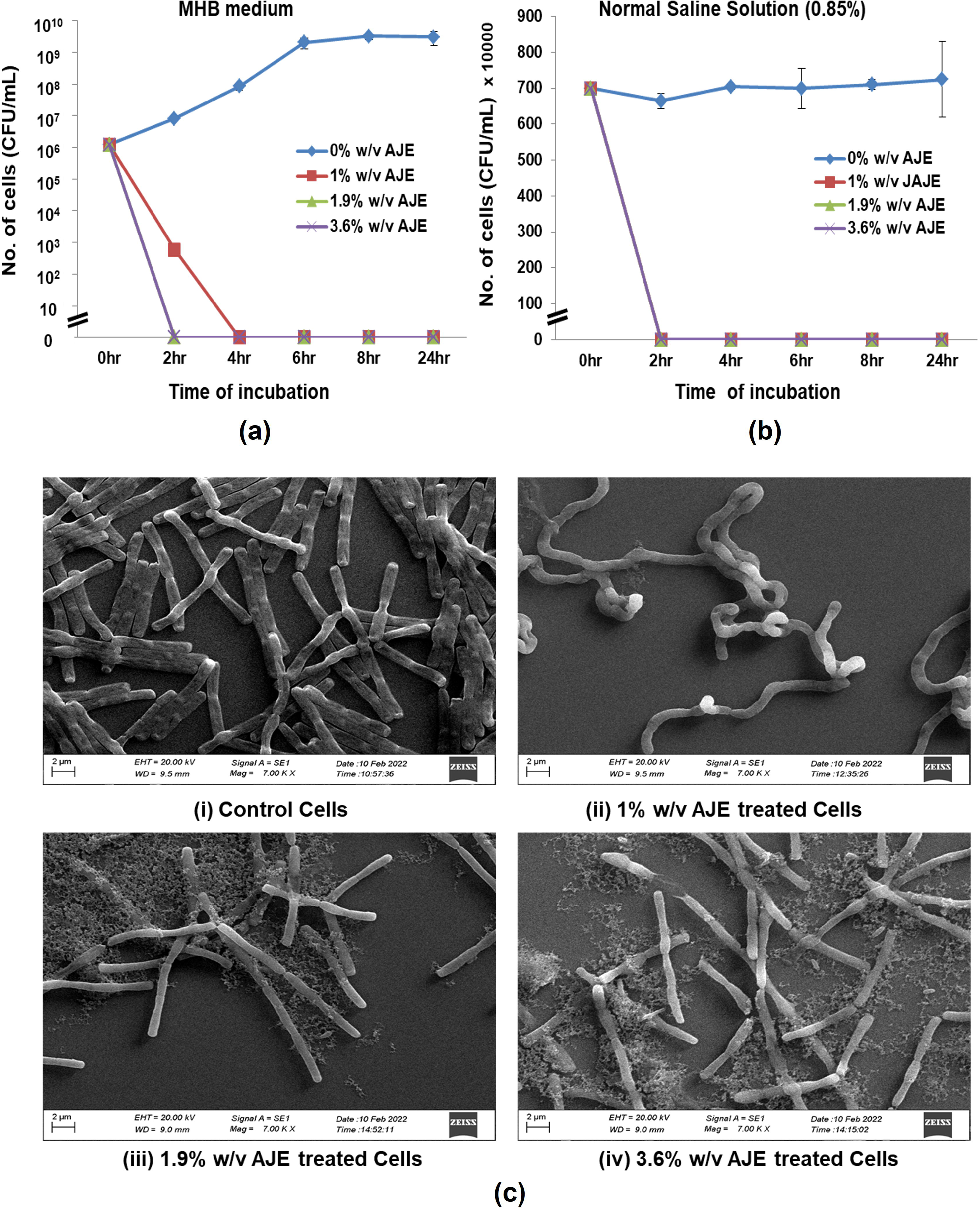
Aqueous Jamun (*Syzygium cumin L.*) extract (AJE) inhibits growth of *B. anthracis*. The exponentially growing *B. anthracis* culture was diluted so as to have 1X 10^6^ CFU/mL, supplemented with indicated concentrations of AJE and growth was monitored both in the presence of organic growth medium, *i.e.*, MHB **(a)**, and absence of organic media, *i.e*., Normal saline solution or NSS **(b)** by enumerating the colony forming units (CFU) on plating of the culture on MHA at different time intervals. The data presented is average of three experiments (n = 3). Error bars represent standard error of the mean (S.E.M). **(c)** Scanning electron microscopy (SEM) image of the exponentially growing *B. anthracis* cells exposed to 0-3.6% w/v AJE for 2 h at 37°C in MHB medium (i-iv). As compared to control cells (i), longer and coiled bacilli chains appeared in 1% AJE exposed cells (ii), while progressive disintegration of longer chains, change in the surface morphology and appearance of cell debris were observed in 1.9% and 3.6% AJE exposed samples (iii-iv).

### 3.4. AJE induces morphological changes in *B. anthracis*

To assess the possible mechanism of AJE’s anti-*B. anthracis* activity, the morphological changes in the *B. anthracis* cells growing in cultures supplemented with 0-3.6% w/v AJE for up to 2 h were examined using Scanning Electron Microscopy (SEM). The control culture of *B. anthracis* that was not exposed to AJE displayed characteristic wild-type morphology, i.e., bacilli growing in chains as expected (Fig. 2c–i). The exposure of growing *B. anthracis* cultures to 1% w/v of AJE is causing the appearance of twisted filamentous morphology, potentially indicative of defective deposition of the cell wall material. Still, exposure to higher concentrations, *i.e.*, 1.9% and 3.6% w/v of AJE, a growth inhibitory concentration, caused elongation of cells, shortening of bacilli chains, and a progressive increase in low-density (white) regions, along with the appearance of extraneous material (debris) [Fig. 2c ii-iv)] within 2 h of AJE exposure. Together, it could be indicative of potential disruption of cell wall synthesis and/or dissolution, along with a loss in viability from AJE exposure.

### 3.5. AJE exposure does not induce virulence plasmid pXO1 loss in *B. anthracis*

The colonies of *B. anthracis* obtained on plating the broth culture grown in the presence of a sub-inhibitory concentration of AJE, *i.e.,* 1% w/v for different time intervals (*i.e.,* 0– 24 h), were found to retain both the pXO1 plasmid-borne *pagA* gene as well as the chromosomal *phoP* gene (Suppl. Fig. 2). All random colonies (n = 6–10) tested displayed amplification of both genes, confirming non-curing of pXO1 from *B. anthracis* at tested sublethal AGE exposure. Therefore, it may be surmised that sublethal concentrations of AJE may not induce virulence plasmid loss in *B. anthracis* (Suppl. Fig. 2).

### 3.6. Anti-*B. anthracis* activity of AJE is stable at high temperature

The comparative ability of AJE and AGE to tolerate different temperatures was evaluated to assess their suitability for storage and usage in different field conditions. The ZOI given by the AJE and AGE after incubation at different temperatures (*i.e.*, 4–50°C) for different time durations (*i.e.,* 0–20 days), indicated that the anti-*B. anthracis* activity of AJE and AGE is relatively stable at 40–50°C (Fig. 3a-b). The AJE retained >90% of its residual anti-*B. anthracis* activity after incubation at 50°C for 20 days, whereas the AGE lost about 50% of its anti-*B. anthracis* activity after 3 days of incubation at 50°C. When the thermal stability of both AJE and AGE was compared at extreme temperatures, *i.e.,* 95°C for 0-12 h it was observed that AJE retained about 80% of its residual anti-*B. anthracis* activity up to 12 h of incubation, whereas AGE lost it completely within 20 minutes of incubation (Fig. 3c). It indicates that the anti-*B. anthracis* activity of AJE is more thermostable than that of AGE.

**Figure 3.**
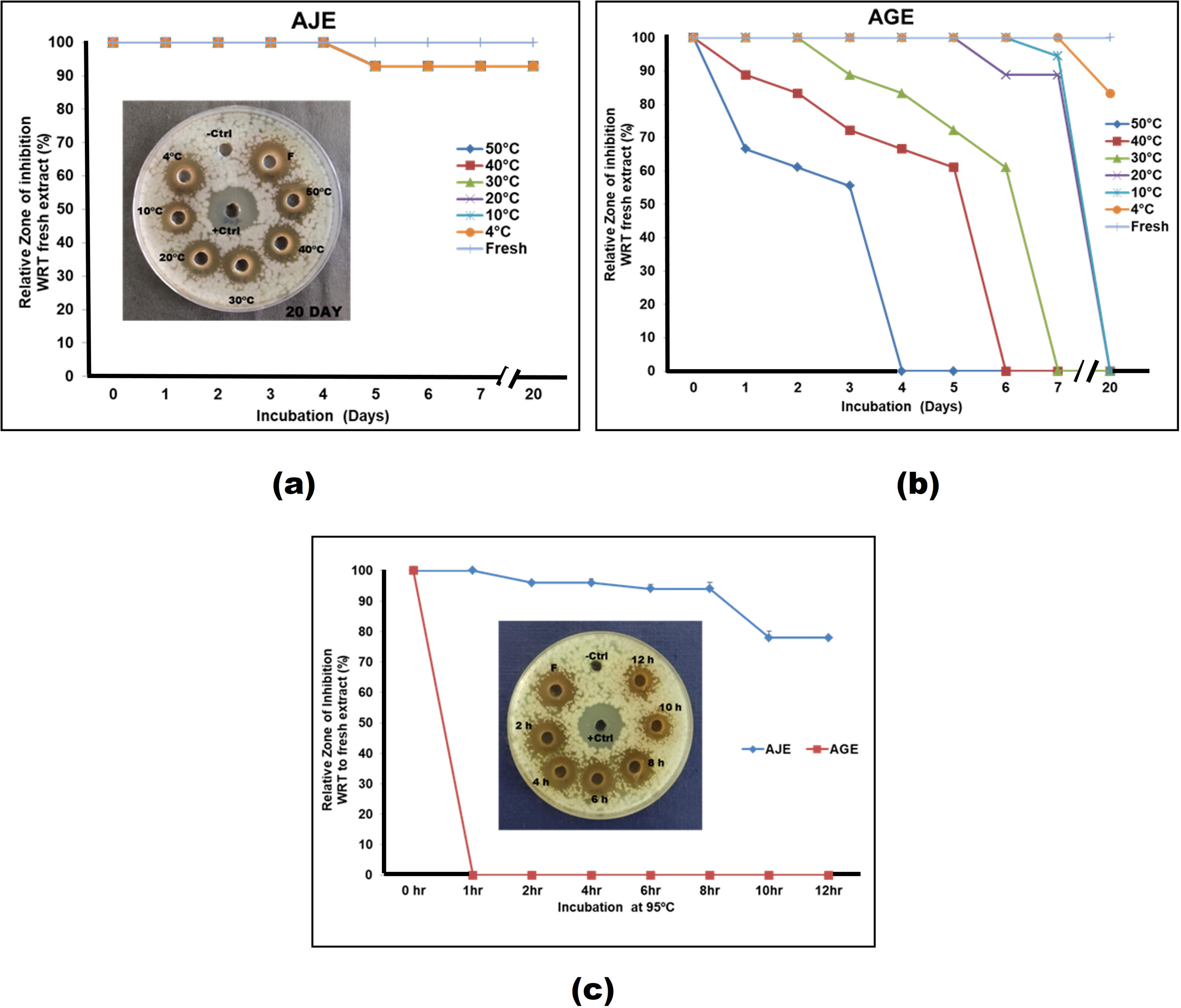
Thermal tolerance of aqueous Jamun Extract (AJE). Exposure of AJE (80% w/v) and AGE (40% w/v) to temperatures up to 95°C does not drastically affect the anti-*B. anthracis* activity of AJE**. (a-b)** The graphs display the relative ZOI as produced by AJE and AGE samples, which were incubated for up to 20 days at temperatures of 4– 50°C. The data is presented as ZOI with respect to (WRT) that produced by fresh AJE. AJE appeared to be quite stable at high temperatures. The AJE incubated at 50°C retained >90% activity even after incubation for 20 days. **(c)** Exposure of AGE and AJE to 95°C decreased the anti-*B. anthracis* activity of AGE but did not seem to affect the activity of AJE. The values reported are the average of three experiments (n = 3). Rifampicin was used as a positive control (+Ctrl) and ultrapure water as a negative control (-Ctrl). A representative image of AWDAs performed to assess the anti-*B. anthracis* activity remaining in AJE (50 μl) after incubation at 0–50 °C and 95 °C is included as an inset in panels (a) and (c).

### 3.7. Different batches of *Syzygium cumini* L. Skeels leaves show similar anti-*B. anthracis* activity

The bioactive components of plants vary with their place of origin, their age, atmosphere, *etc.* due to which there can be variability in their antibacterial activity. The shown representative extracts from three different leaf samples from different locations in Varanasi display quite similar anti-*B. anthracis* activity forming 14–15 mm size ZOIs (Suppl. Fig. 3). The results suggest anti-*B. anthracis* activity in AJE leaves is unaffected by location.

### 3.8. Anti-*B. anthracis* activity in AJE is relatively stable in the physiologically relevant conditions and displays selectivity towards *B. anthracis* but is harmless for other gut microbiota

The ability of AJE to tolerate different physiological conditions present in the host gut such as bile salts, gut microbes, and extreme pH was tested to assess its potential application in anthrax treatment and control. The presence of bile salts [*i.e.,* sodium deoxycholate (0.0025% w/v)] did not antagonize AJE anti-*B. anthracis* activity. The antibacterial activity of AJE against *B. anthracis* and *E. coli* DH5α indicated that *B. anthracis* cells are much more sensitive to AJE than *E. coli* (Fig 4a-i and 4b). The comparison of the antibacterial activity of AJE and AGE against *B. anthracis* and other gut microbes present in two commercially available probiotic formulations indicated that *B. anthracis* cells are highly sensitive to AJE exposure as compared to other gut microbes, unlike AGE that displays much lower discriminatory activity towards other gut microbes present in probiotics, indicative of significantly lower selectivity (Fig. 4b). Thus, AJE displays selectivity towards *B. anthracis*, a pathogen, which is a property desired in any ideal antimicrobial agent.

**Figure 4.**
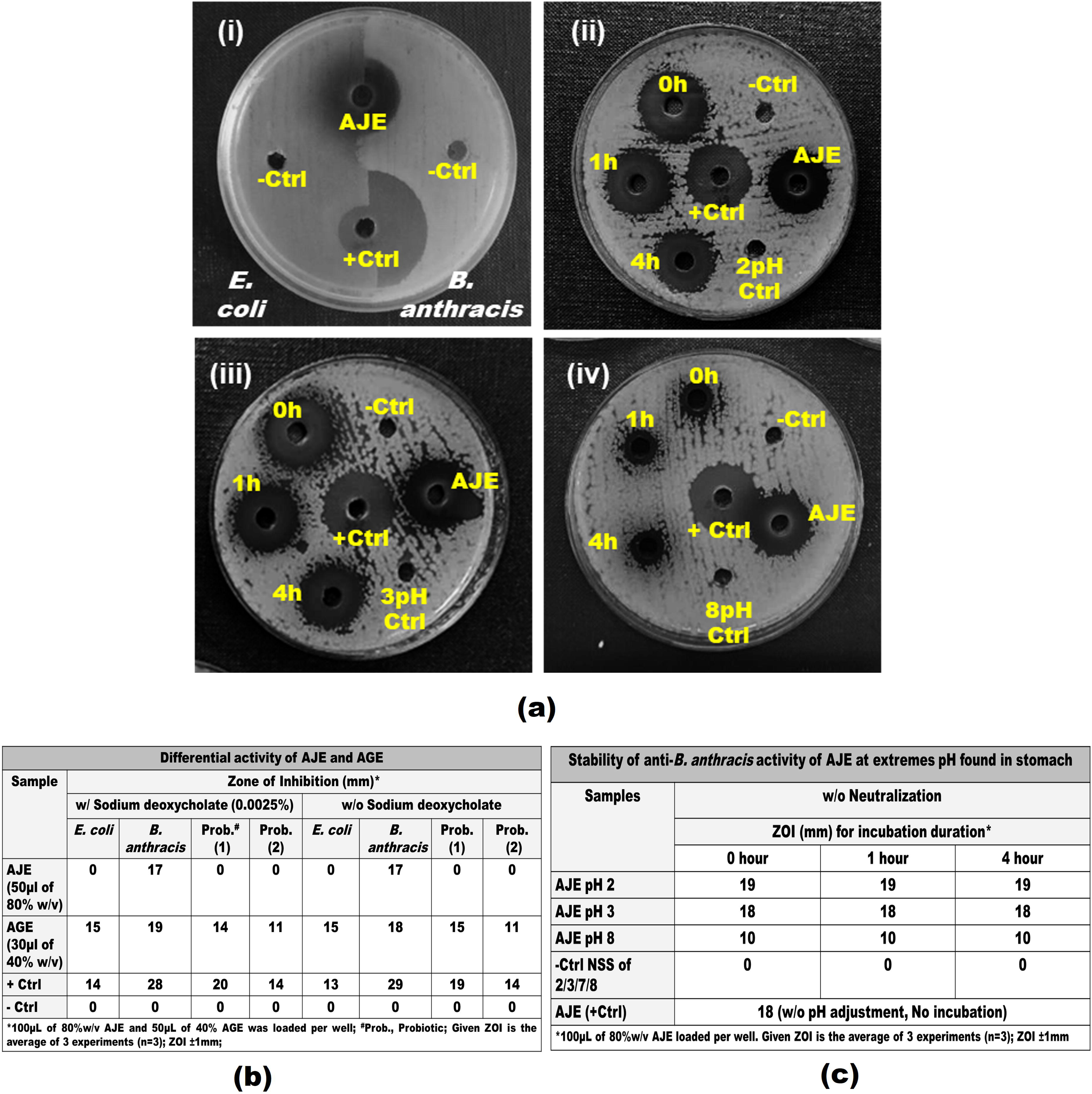
Characterization of Aqueous Jamun (*Syzygium cumini* L.) Extract (AJE). (**a-i**) A representative image of AWDA performed to compare the relative sensitivity of the *Escherichia coli* DH5α strain and the *B. anthracis* Sterne strain to AJE, head-to-head, on a MHA plate supplemented with sodium deoxycholate (0.0025%). [Likewise, AWDA was performed for the relative sensitivity assessment of microbes commonly present in probiotics; refer to Panel (b) for comparison]. **(a-ii-iv)** The effect of different pH conditions (*i.e.,* 2, 3, and 8) on AJE anti-*B. anthracis* activity: Aliquots of pH-adjusted AJE (*i.e.,* pH 2, 3, and 8) were incubated for 0, 1, and 4 h at 37°C. The 100μl aliquots of the incubated samples were assessed for the remaining or residual anti-*B. anthracis* activity by AWDA. (-Ctrl/ 2pH/ 3pH/8pH: saline/pH adjusted saline; +Ctrl: Rifampicin). **(b-c)** Summary of the assay shown in (a; i–iv): The ZOI of AJE may be compared with that of AGE. It shows that AJE does not inhibit the growth of *E. coli* and other gut microbes present in the tested probiotics, irrespective of the presence of the bile salt sodium deoxycholate (0.0025%). Probiotic (1) and Probiotic (2) are commercially available probiotics with different compositions of microbes (see Section 2.7 for the identity of microbes). Together, these findings suggest that the use of AJE as a therapeutic agent would potentially neither disturb the microflora of the host intestine nor cause a loss in its activity at the acidic pH encountered in the stomach, making it a better anti-*B. anthracis* candidate than AGE.

The relative ZOI produced by pH adjusted AJE incubated for 0, 1, and 4 h at 37°C indicates that the anti-*B. anthracis* activity of AJE is quite stable at pH 2 and 3 [Fig 4a (ii-iv) and 4c]. At pH 8, it was observed that anti-*B anthracis* activity of AJE gets reduced but this decrease is reversible as when pH 8 of the AJE is again adjusted to pH 2 or 3, its anti-*B. anthracis* activity is restored (Suppl. Fig. 4). (Note: It was observed that at pH 8, AJE precipitates and when again its pH is adjusted to pH 2 or 3, precipitates disappear – go back to solution)

### 3.9. AJE does not antagonize the activity of antibiotics recommended for *B. anthracis* treatment

The FDA recommended antibiotics prescribed for anthrax treatment such as Amoxicillin, Cefixime, Ciprofloxacin, Doxycycline, Levofloxacin, Penicillin, Rifampicin, and Tetracycline (CDC, 2016; U.S. Food and Drug Administration, 2018) were tested for their possible interaction with AJE. As expected, all the approved antibiotics inhibited the growth of *B. anthracis.* The AJE did not seem to antagonize the anti-*B. anthracis* activity of any of the tested antibiotics recommended for the treatment of anthrax. Further experiments indicated a synergistic interaction of AJE with Amoxicillin, Rifampicin, and Tetracycline (FICI<1), whereas an indifferent interaction was displayed with Ciprofloxacin, Doxycycline, Levofloxacin, and Penicillin (Fig. 5). It can be concluded from the observation that AJE can be used along with the recommended antibiotics for anthrax control.

**Figure 5.**
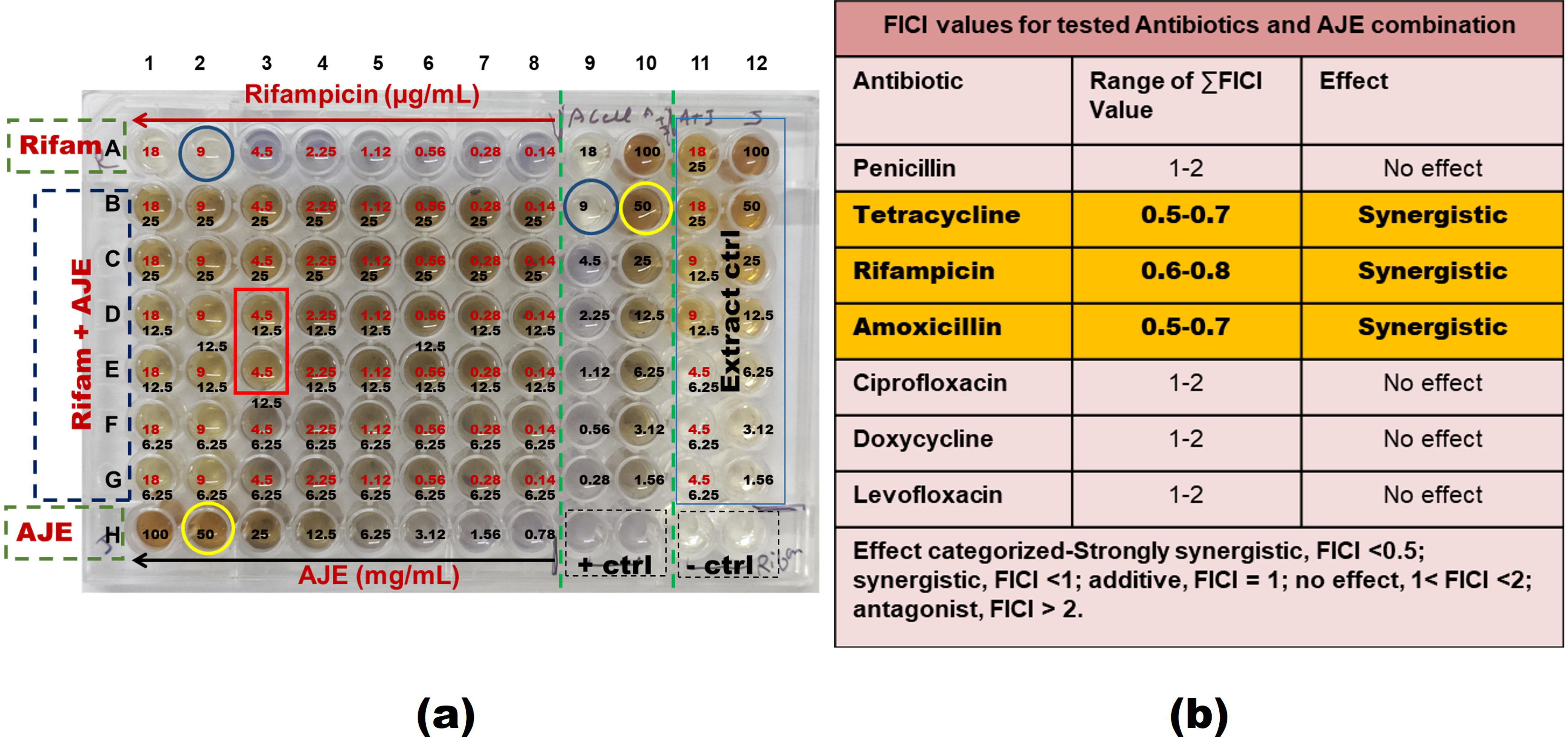
Aqueous Jamun Extract (AJE) does not antagonize the anti-*B. anthracis* activity of the commonly employed antibiotics for anthrax control. (**a**) A typical microdilution experiment for MIC and FICI value calculation is shown. A representative plate showing interaction between Rifampicin (Rifam) (*i.e.,* 18–0.14 μg/ml; red) and AJE (*i.e.,* 100–0.78 mg/mL; black) is displayed. The concentration of Rifampicin and AJE alone decreased from left to right (row A and H), while the concentration of AJE in combination with the indicated fixed concentration of antibiotic increased from bottom to top (row G to B) as indicated by the arrows. Column 9 and 10 are the duplicates of row A and H, respectively. The wells H11-12 are media control (-Ctrl) while H9-10 are cell control, *i.e., B. anthracis* cells without any antibiotics or AJE. Values indicated in the blue circle are MIC value of Rifampicin (*i.e.,* 9 μg/ml) and that in yellow circle are MIC value of AJE (*i.e.,* 25 mg/mL or 2.5% w/v) in the rows A and H, respectively. Values in the red box indicate the concentration of Rifampicin and AJE in combination (*i.e.,* 4.5 μg/ml and 12.5 mg/ml respectively). **(b)** Fractional inhibitory concentration index (FICI) calculation for the antibiotics and AJE combinations. The values represent the range of three experiments performed independently. The concentration range of antibiotics used in combination with AJE was as follows: Amoxicillin (6–0.02 μg/ml), Tetracycline (2– 0.15 μg/ml), Ciprofloxacin (2–0.15 μg/ml), Levofloxacin (2.5–0.19 μg/ml), Doxycycline (2-0.15 μg/ml) and Penicillin (50-0.39 μg/ml).

### 3.10. TLC-bioautography of AJE and Mass spectrometry analysis of bioactive fraction

The separation of bioactive fractions with anti-*B. anthracis* activity from crude AJE was done by the TLC method followed by bioautography to identify the bioactive fraction/spots (Suppl. Fig. 5a). The separated bioactive fraction from the area corresponding to the anti-*B. anthracis* activity on the duplicate TLC plate in bioautography, was scratched from the plate and subjected to GC-MS to detect the potential bioactive fraction/compound. The bioautography indicated spots where AJE was loaded to have anti-*B. anthracis* activity (Suppl. Fig. 5b); encircled in red in the right panel: AJE TLC plate. The area on TLC plates where AJE was fractionated but no anti-*B. anthracis* activity was observed, was used as a negative control for comparison (labeled yellow circle, Suppl. Fig. 5b). Both spots with and without anti-*B. anthracis* activity was scratched from the TLC plate and GC-MS was performed. The GC-MS profile/outline for both the spots with and without anti-*B. anthracis* activity is shown in Suppl. Fig. 5c-d. The potential list of the bioactive compound ascertained from the NIST 2.0 (National Institute of Standards and Technology) library is listed in Table 3. The anti-*B. anthracis* activity displaying bioactive fraction constituents of AJE included various bioactive compounds of different classes belonging to categories of alkaloids, glycosides, tannins, saponins, terpenes, steroids, flavonoids, and phenols with potential antibacterial activity. Some of the known antimicrobial compounds of different classes are identified in GC-MS analysis of AJE (Table 3) such as Phytol (Cyclic diterpene), 9-Octadecenamide (Fatty acid), Dodecanamide (alky amides), 1,3-Dioxolane (Ketones), 1-Decanol, 5,9-dimethyl and 3-Pentadecanol (long-chain fatty alcohols), Octadecane, 1-ethenyloxy (terpenoids). Many other compounds with anti-inflammatory, anti-pruritic, anti-diabetic, anticancer, antioxidant, anti-allergic, or vasoconstrictive actions were also identified from the bioactive fraction. The compounds that cover the highest peak in the GC-MS, like 2-Acetoxyisobutyryl chloride, 1,3-Dioxolane, 2,2,4-trimethyl, and 1,3-Dioxolane-2-methanol, 2,4-dimethyl were already reported to have antibacterial and antifungal activity (See Table 3). Thus, the characterization of the bioactive fraction indicates that the observed anti-*B. anthracis* activity of the AJE could be due to one or more antimicrobial compounds.

**Table 3:** List of compounds identified from AJE Bioactive fraction by GC-MS.

| S. No. | Compounds identified | Mol. Formula | RT | Area | Properties | Reference |
| --- | --- | --- | --- | --- | --- | --- |
| 1. | 2-Acetoxyisobutyryl chloride | C<sub>6</sub>H<sub>9</sub>ClO<sub>3</sub> | 3.6 | 63 | Antibacterial<br>Antifungal | Dr. Duke and NIST<br>Phytochemical and<br>Ethnobotanical<br>databases |
| 2. | Propane, 1,1-dipropoxy | C<sub>9</sub>H<sub>20</sub>O<sub>2</sub> | 3.6 |  | No activity reported | - |
| 3. | 1,3-Dioxolane, 2,2,4-trimethyl | C<sub>6</sub>H<sub>12</sub>O<sub>2</sub> | 3.6 |  | Antibacterial<br>Antifungal | Kucuk *et al.*, 2011 |
| 4. | 1,3-Dioxolane-2-methanol, 2,4-dimethyl | C<sub>6</sub>H<sub>12</sub>O<sub>3</sub> | 3.6 |  | Antibacterial<br>Antifungal | Kucuk *et al.*, 2011 |
| 5. | Diazene, dimethyl | C<sub>2</sub>H<sub>6</sub>N<sub>2</sub> | 3.96 | 8.18 | Antitumor,<br>Antimicrobial<br>Antibacterial | Dembitsky *et al.*, 2017 |
| 6. | Tetramethyl ammonium acetate | C<sub>6</sub>H<sub>15</sub>NO<sub>2</sub> | 3.96 |  | No activity reported | - |
| 7. | Acetamide, n-propyl-2-(4H-[1,2,4]triazol-3-yl) | C<sub>7</sub>H<sub>12</sub>N<sub>4</sub>O | 7.94 | 0.13 | Antiviral | Kharb *et al.*, 2011 |
| 8. | Oxalic acid, cyclohexyl propyl ester | C<sub>11</sub>H<sub>18</sub>O<sub>4</sub> | 7.94 |  | No activity reported | - |
| 9. | Oxalic acid, cyclohexyl pentyl ester | C<sub>13</sub>H<sub>22</sub>O<sub>4</sub> | 7.94 |  | No activity reported | - |
| 10. | Oxirane, tetramethyl | C<sub>6</sub>H<sub>12</sub>O | 8.71 | 0.28 | Antimicrobial | Thirunarayanan 2014 |
| 11. | 3-Hexanol, 4-ethyl | C<sub>8</sub>H<sub>18</sub>O | 8.71 |  | No activity reported | - |
| 12. | (2,3,3-Trimethyloxiranyl) methanol | C<sub>6</sub>H<sub>12</sub>O<sub>2</sub> | 8.71 |  | No activity reported | - |
| 13. | 7-Nonenamide | C<sub>9</sub>H<sub>17</sub>NO | 15.54 | 3.7 | No activity reported | - |
| 14. | Cyclohexane carboxamide | C<sub>7</sub>H<sub>13</sub>NO | 15.54 |  | Antibacterial<br>Anti-yeast | Arslan *et al.*, 2009 |
| 15. | 3-Cyclohexylpropionamide | C<sub>9</sub>H<sub>17</sub>NO | 15.54 |  | Antimicrobial | Temiz-Arpaci *et al.*, 2002 |
| 16. | N'-(4-Nitrobenzylidene)-2-pentyl-1-cyclopropanecarbohydrazide | C<sub>16</sub>H<sub>21</sub>N<sub>3</sub>O<sub>3</sub> | 15.77 | 2.42 | No activity reported | - |
| 17. | 1-Decanol, 5,9-dimethyl | C<sub>12</sub>H<sub>26</sub>O | 15.77 |  | Antibacterial | Mukherjee *et al.*, 2013 |
| 18. | 9-Octadecenamide, (Z) | C<sub>18</sub>H<sub>35</sub>NO | 16.04 | 2.6 | Hepatoprotective<br>Antihistaminic<br>Aypocholesterolemic<br>Antieczemic | Dr. Duke and NIST<br>Phytochemical and<br>Ethnobotanical<br>databases |

**Table 3:** List of compound identified from AJE Bioactive fraction by GC-MS
|  |  |  |  |  |  |  |
| --- | --- | --- | --- | --- | --- | --- |
| 19. | Dodecanamide | C<sub>12</sub>H<sub>25</sub>NO | 16.04 |  | Antibacterial<br>Antifungal | Dr Duke and NIST<br>Phytochemical and<br>Ethnobotanical<br>databases |
| 20. | 3-Pentadecanol | C<sub>15</sub>H<sub>32</sub>O | 16.59 | 0.22 | Antibacterial | Togashi *et al.*, 2007 |
| 21. | 3-Heptadecanol | C<sub>17</sub>H<sub>36</sub>O | 16.59 |  | Antibacterial | Togashi *et al.*, 2007 |
| 22. | Phytol | C<sub>20</sub>H<sub>40</sub>O | 19.42 | 0.07 | Antimicrobial<br>Anticancer<br>Antioxidant<br>Diuretic<br>Anti-inflammatory | Willie *et al.*, 2021 |
| 23. | Octadecane, 1-(ethenyloxy)- | C<sub>20</sub>H<sub>40</sub>O | 19.42 |  | Antibacterial<br>Antifungal | Naqvi *et al.*, 2019 |
| 24. | 1-Dodecanol, 3,7,11-trimethyl | C<sub>15</sub>H<sub>32</sub>O | 19.42 |  | No activity reported | - |
| 25. | 9-Octadecenoic acid (Z)-, phenylmethyl ester or benzyl oleate | C<sub>25</sub>H<sub>40</sub>O<sub>2</sub> | 25.22 | 0.21 | No activity reported | - |
| 26. | Octadec-9-enoic acid | C<sub>18</sub>H<sub>34</sub>O<sub>2</sub> | 25.22 |  | Antimicrobial | M. M. Rahman *et al.*, 2014 |
| 27. | 9-Octadecenoic acid (Z)-, 2-hydroxyethyl ester | C<sub>20</sub>H<sub>38</sub>O<sub>3</sub> | 25.22 |  | Antioxidant<br>Anti-inflammatory | Duke 2008 |
| 28. | 6-Octadecenoic acid | C<sub>18</sub>H<sub>34</sub>O<sub>2</sub> | 25.22 |  | Anti-inflammatory<br>Anti-androgenic Anti-cancer<br>Hypocholesterolemic | Sreekumar *et al.*, 2014 |

## 4. DISCUSSION

Anthrax is an important endemic disease in many developing nations. In the countries of Africa and Asia, anthrax is a matter of greater concern as they share a larger livestock population and relatively poor veterinary healthcare systems (Kaur et al., 2013; Manish et al., 2020; Shadomy et al., 2016; Turnbull, 2008). There is a significant impact of zoonotic diseases on public health, animal economies, and wildlife in India. Anthrax remains one of the most significant zoonotic diseases in parts of India (Bhattacharya et al., 2021). With the global rise in antimicrobial resistance among pathogens (Singh et al., 2022), there is a growing need for the discovery of novel antibacterial agents with new or multiple modes of action to effectively reduce the pathogenicity of organisms by controlling their multiplication or division (Bhardwaj et al., 2013; Ta and Arnason, 2016) or causing the loss of virulence plasmids (*e.g*., pXO1, pXO2) so that the spread and infections can be effectively controlled, even in the case of toxemia (Dastidar et al., 2013; Spengler et al., 2006).

From ancient times plants are considered an important source of antimicrobial compounds which are known for controlling various diseases caused by pathogenic microorganisms (Dastidar et al., 2013; Mahady, 2005; Ta and Arnason, 2016). The identification of such livestock edible plants or their parts that can be included in their feed and fodder and has anti-*B. anthracis* activity too could directly impact the prevention and treatment of anthrax or anthrax-like GI infections among livestock in poorer endemic regions. The organic and aqueous extracts obtained from different plant parts have been reported in the literature to have anti-*B. anthracis* activity (Akinpelu et al., 2008; Elisha et al., 2016; Kaur et al., 2021; Mbwambo et al., 2011; Taher et al., 2012). In our study, several livestock edible plants commonly found in the Indian subcontinent (Duke, 2008, 2002; USDA, 2016) that are reported to have antimicrobial or stomachic properties and indicated for dysentery (diarrhea or bloody diarrhea), stomachache, bloating of the stomach, vomiting, fever and chills, etc., were evaluated for anti-*B. anthracis* activity. We have selected water (aqua) as an extractant for its ease of potential field applicability, unlike other organic solvents. Among the tested aqueous plant extracts, the aqueous *Syzygium cumini* (Jamun) extract (AJE) was found to have the most potent anti-*B. anthracis* activity that consistently inhibited the growth of *B. anthracis*. However, its relative potency was lower than that of previously reported aqueous garlic extract (AGE) (Kaur et al., 2021). In literature, Jamun has been also reported for its therapeutic and herbal properties as anti-hyperglycemic, anti-inflammatory, antimicrobial, cardioprotective, anti-diabetic diarrhea, bilious diarrhea, appetite increasing, anti-dysentery, etc. (Arun et al., 2011; do Nascimento-Silva et al., 2022; Duke, 2008, 2002; Jagetia, 2018; Rekha et al., 2008; Sharma et al., 2008; Tanwar et al., 2011; Yadav et al., 2017). It is indicated to play an important role in the prevention and management of non-communicable diseases such as diabetes mellitus, cancer, gout, heart disease, *etc.,* as well (Ahmad et al., 2019). Previously, the aqueous extract of Jamun stem and leaf had been shown to have antibacterial activity against *Staphylococcus aureus, Staphylococcus saprophyticus, Escherichia coli, Pseudomonas aeruginosa,* and *Proteus vulgaris*, whereas the fruit extracts were active against *P. aeruginosa* and antifungal activity against *Penicillium chrysogenum* and *Candida albicans* (Jagetia, 2017). However, we did not observe anti-*E. coli* activity in AJE at the tested concentration. We speculate, possibly higher concentration may inhibit *E. coli* growth. The extracts of *Syzygium cumini* leaf had been also reported to inhibit the biofilm formation and virulence of *S. aureus* and *P. aeruginosa* (Gupta et al., 2019). In our study, at the sub lethal concentration of AJE, *i.e*., 1% w/v the *B. anthracis* cells developed coiled or twisted asymmetric shape/structure. We speculate this could be due to the changes in cross-linked peptidoglycan (PG) structure in the cell wall as proposed previously by Yang et al., 2016. The predicted mechanism of action is shown in Suppl. Fig. 6.

The toxins produced by vegetative cells of *B. anthracis* are the main cause of its pathogenesis (Omotade et al., 2014). Though AJE at higher concentrations displayed efficient killing of *B. anthracis* cells, the ability to promote/cause virulence plasmid loss at tested sublethal concentration was not observed, similar to that was previously observed for the aqueous Garlic extracts (Kaur et al., 2021).

Identification or characterization of bioactive fractions/compounds from crude plant extracts is required to correlate the activity of the individual compound and their effects on pathogenic microorganisms so that its proper preventive or curative use in controlling diseases can be recommended (Bhardwaj et al., 2013; Friedman, 2015; Guil-Guerrero et al., 2016; Elkenawy et al., 2023). Sagrawat et al., (2006) had suggested that the leaves of *Syzygium cumini* L. Skeels contain various chemicals such as β-sitosterol, betulinic acid, mycaminose, crategolic (maslinic) acid, *etc.,* which have medicinal value (Sagrawat, 2006). Eshwarappa et al. have indicated the methanolic and aqueous leaf extract of Jamun to contains various phytochemicals like phenols, tannin, flavonoids, phytosterols, triterponoids, alkaloids, and saponins with antioxidant and antimicrobial properties which could be bioactive principles behind their indication for treatment of various diseases like diabetes mellitus, arthritis, cancer, liver disorder, *etc.*, (Eshwarappa et al., 2014). Jamun has been reported to contain various phytochemicals like alkaloids, catechins, flavonoids, glycosides, steroids, phenols, tannins, saponins, anthraquinones, and cardiac glycosides (Jagetia, 2018). Our analysis of the AJE indicated the presence of these classes of compounds. The compounds identified by GC-MS analysis with the maximum coverage, like 2-Acetoxyisobutyryl chloride and Diazene dimethyl, already reported in the literature for their antimicrobial activity (Duke 2008; Dembitsky et al., 2017), could be responsible for the anti-*B. anthracis* activity displayed by AJE. The potential contribution/role of other compounds present in smaller quantities cannot be ignored. However, future investigations would provide a more definitive answer to their potential application in controlling anthrax.

When one or more antimicrobial agents or drugs are used in combination, they may interact (synergistically or antagonistically) or remain indifferent (Acar, 2000; Weiss et al., 2015; Wolfart et al., 2006). Whenever a new antimicrobial agent has to be introduced, this kind of information (its interaction with other agents, *i.e.,* synergistic, antagonistic, or indifferent) is needed for its successful usage in combination with the known in-use antibacterial agent. In our study, the AJE displayed safer interaction with available and recommended antibacterial agents as it did not seem to antagonize the anti-*B. anthracis* activity of any of the tested FDA-approved antibiotics currently used for anthrax control. Therefore, the usage of AJE along with the combination of other FDA-approved antibiotics to reduce anthrax or anthrax-like infections could not potentially pose a problem.

The thermal stability characterization of AGE indicated that the anti-*B. anthracis* activity of AJE constituents is quite stable at high temperatures (40–50°C). It retained more than 90% of its anti-*B. anthracis* activity for 20 days at 50°C and at 95°C it retained 80% of its activity up to 12 h, whereas in the case of aqueous garlic extract AGE, the anti-*B. anthracis* activity was lost within 2 hrs. The anti-*B. anthracis* activity of Jamun leaves could even tolerate boiling, as the aqueous extract of leaves obtained on boiling (10 min) retained >95% of its anti-*B. anthracis* activity (data not shown). This indicates that AJE has quite thermostable anti-*B. anthracis* activity, which could allow its wider application in the field for treating anthrax and anthrax-like GI diseases. The increased thermal stability of AJE makes it a better potential candidate than AGE for therapeutic usage in the future.

The anti-*B. anthracis* activity of AJE was found to be quite stable at the acidic pH (*i.e.,* 2-3 pH) encountered in the host intestine. Near neutral/basic pH (*i.e.,* 7-8 pH) the AJE shows precipitation and reduction in anti-*B. anthracis* activity but once its pH is adjusted to its original (∼4 pH) or at 2 or 3 pH the precipitates dissolve and the anti-*B. anthracis* activity of AJE is retrieved, indicating a non-destructive reduction in activity on pH change.

The commensal *E. coli* and other gut microbes (commonly present in the gut of the host, also taken as probiotics (*e.g., Streptococcus faecalis, B. mesentericus, Clostridium butyricum, Lactobacillus* spp*., Streptococcus thermophiles, Saccharomyces boulardii, Bifidobacterium* spp.) were found to be resistant to AJE in the presence and absence of sodium deoxycholate, unlike *B. anthracis*. This indicates that the original microflora of host intestines would not get disturbed by AJE consumption, moreover, bile salts (sodium deoxycholate) present in the host intestine are presumably not going to interfere with AJE’s anti-*B. anthracis* activity. In our previous study, we reported that *E. coli* is sensitive to AGE (Kaur et al., 2021). In the current study, our comparison of the sensitivity of *E. coli* and a number of gut commensals present in the two probiotic formulations to AGE and AJE, indicates that the *E. coli* and the microbes present in the tested probiotics have better tolerance to AJE as compared to AGE. The relative resistance of *E. coli* and other gut commensals (present in probiotics) to AJE constituents, in contrast to *B. anthracis,* could be supposedly a result of their coevolution with hosts, where they were frequently exposed to edible plants such as *Syzygium cumini* L. Skeels (Jamun) as fodder or feed. Furthermore, genetic exchange mechanisms could have also played a role in the possible dissemination of the tolerance genes among them once acquired by a commensal during the coevolution.

Further work needs to be undertaken to explore the possibility of employing *Syzygium cumini* (Jamun) leaf or its extract to decrease the incidences of anthrax in endemic areas. Immediate future work may be focused on determining the mode of action of AJE and purifying the compound(s) responsible for the anti-*B. anthracis* activity and testing it in animal models. The assessment of AJE as an oral or an injectable for therapy would be an area of exploration as well, as seed, bark, fruit, leaf powder and decoctions of Jamun are known to be well tolerated (Duke, 2008; FSSAI, 2016; Ministry of Ayush, 2016).

## 5. CONCLUSION

The evaluation of the aqueous plant extracts of livestock edible plants, which are traditionally indicated to treat various gastrointestinal (GI) infections including GI anthrax/anthrax-like diseases suggested that the aqueous extract of *Syzygium cumini* L. Skeels (AJE) has potent anti-*B. anthracis* activity. The aqueous Jamun (*Syzygium cumini* L. Skeels) extract (AJE) has greater stability under different assayed conditions that are relevant to its use in anthrax treatment and control, such as its exposure to a higher temperature (up to 95°C), stability at acidic pH conditions, and tolerance to bile salts. Furthermore, AJE does not adversely affect the gut microflora and does not antagonize the activity of FDA-approved antibiotics recommended for anthrax treatment indicating its compatibility with available therapeutic medication and treatments for anthrax. The identification and characterization of the bioactive fraction constituents indicated the presence of various derivatives of acid esters, phthalic acid, steroids and phenyl group containing compounds with potential antimicrobial activity. Further characterization of the bioactive compounds present in the AJE and their mode of action(s) could open up new avenues to control anthrax and anthrax-like diseases. Together, these findings support the evidence-based use of *Syzygium cumini* L. Skeels for GI anthrax control in endemic areas.

## AUTHOR CONTRIBUTIONS

Conceived and designed the experiments: SS; Performed the experiments: RK; Analyzed the data: RK, SS; Contributed reagents/ materials/analysis tools/discussion: RK, AKS, IKM and SS.

## ACKNOWLEDGEMENT

The work was supported by ‘Ramalingaswami Fellowship (BT/RLF/Re-Entry/50/2011)’ from the Department of Biotechnology (DBT) and the funding support from the Institute of Eminence (IoE) seed grant, Banaras Hindu University, India to SS; RK is a recipient of the UGC Non-NET fellowship by Banaras Hindu University, Varanasi, India.

## FIGURE LEGENDS

**Supplementary Figure 1: Anti-*B. anthracis* activity of different solvent extracts of *S. cumini* L. Skeels (*i.e.,* Methanol, Chloroform, Acetone, Hexane, Butanol and Ethyl acetate): (a)** AWDA of different solvent extracts of *S. cumini;* (b) Comparative ZOI of different solvent extracts of *S. cumini* L. Skeels (*i.e.,* Methanol, Chloroform, Acetone, Hexane, Butanol and Ethyl acetate). Note: +: solvent with *S. cumini* leaf extract; d: dried extract dissolved in water, +ve; ctrl; Rifampicin used as a positive control (80µg/ml), –ve ctrl: solvents only (only chloroform showed activity among the tested solvents).

**Supplementary Figure 2: Aqueous Jamun Extract (AJE) at sub-inhibitory concentration does not promote loss of virulence plasmid ‘pXO1’ from *B. anthracis* Sterne strain.** The *B. anthracis* cells were grown in MHB medium in the presence of sub-inhibitory concentration of AJE (≤ 1% w/v) and the loss of virulence plasmid pXO1 from cells was assessed at different time intervals (0 – 24 hr). The analysis of random colonies picked from 0 and 24 h AJE exposed culture for the presence of *pagA* **(a)** and *phoP* **(b)** genes are shown. All colonies examined retained the pXO1 plasmid based upon the PCR amplification of *pagA* gene **(c)**. Genomic DNA from *B. anthracis* Sterne strain was used as positive control and that of *E. coli* DH5α strain as negative control for the PCR based analysis.

**Supplementary Figure 3: Comparison of the anti-*B. anthracis* activity of AJE** made from three different batches from BHU, Varanasi, India by AWDA. The ZOI values reported are the average of three experiments (n = 3) performed in duplicate.

**Supplementary Figure 4: Effect of pH change on AJE’s anti-*B. anthracis* activity: (a-b)** AWDA of AJE adjusted to pH 2, 3, and 7. Note the effect of reversing the pH of AJE (*i.e.,* from pH 2 and 3 to pH 7 and then again from pH 7 to 2 and 3, respectively) on its anti-*B. anthracis* activity. The 100 μl aliquots of the incubated samples were assessed for the remaining/residual anti-*B. anthracis* activity by AWDA (W: –Ctrl, *i.e.*, water; +Ctrl: Rifampicin).

**Supplementary Figure 5: Thin layer chromatography (TLC) and bioautography of AJE: (a-b)** Separation of bioactive fraction from AJE by using two solvent system, *i.e.,* (A) Methanol: Chloroform (7:3 ratio); (B) Toluene: Acetone (7:3 ratio); **(a)** Indicates the short UV images of TLC silica plate on which AJE was separated by using selected ‘A’ and ‘B’ solvents; **(b)** Bioautography in which anti-*B. anthracis* activity (Zone of inhibition, ZOI) was observed in case of Toluene: Acetone solvent system (B) at the spot where AJE was loaded (circled in red). Area circled in yellow does not display any anti-*Bacillus anthracis* activity and used as a negative control for GC-MS analysis; **(c)** GC-MS profile of spot where ZOI was observed – circled in red in (b); **(d)** GC-MS profile of spot where no ZOI was observed – circled in yellow in (b). List of compounds identified in bioactive spot (circled in red) by eliminating the compounds identified in non-bioactive area of TLC plate (circled in yellow) are given in Table 2.

**Supplementary Figure 6: Predicted mechanism of action of AJE:** AJE could be disrupting the crosslinking of cell wall constituents. Possibly, at a sub-lethal concentration of AJE (1%), the changes in the monomeric and cross-linked peptidoglycan (PG) structure of the N-acetylglucosamine (G)–N-acetylmuramic acid (M) disaccharide would be affecting the shape of the *B. anthracis* cells. The curved and helical shape of bacteria could be the result of an uneven distribution of cell wall (*i.e.,* PG) synthesis or modification. Specifically, areas with asymmetric wedges or zones of higher PG formation would experience increased growth on that side, leading to curvature. Conversely, areas with less PG formation will result in a bend in the opposite direction.

