## Supplementary figures and images for "*Syzygium cumini* (L.) Skeels has Thermostable and Selective Anti-*Bacillus anthracis* Activity"

### Suppl. Figure 1

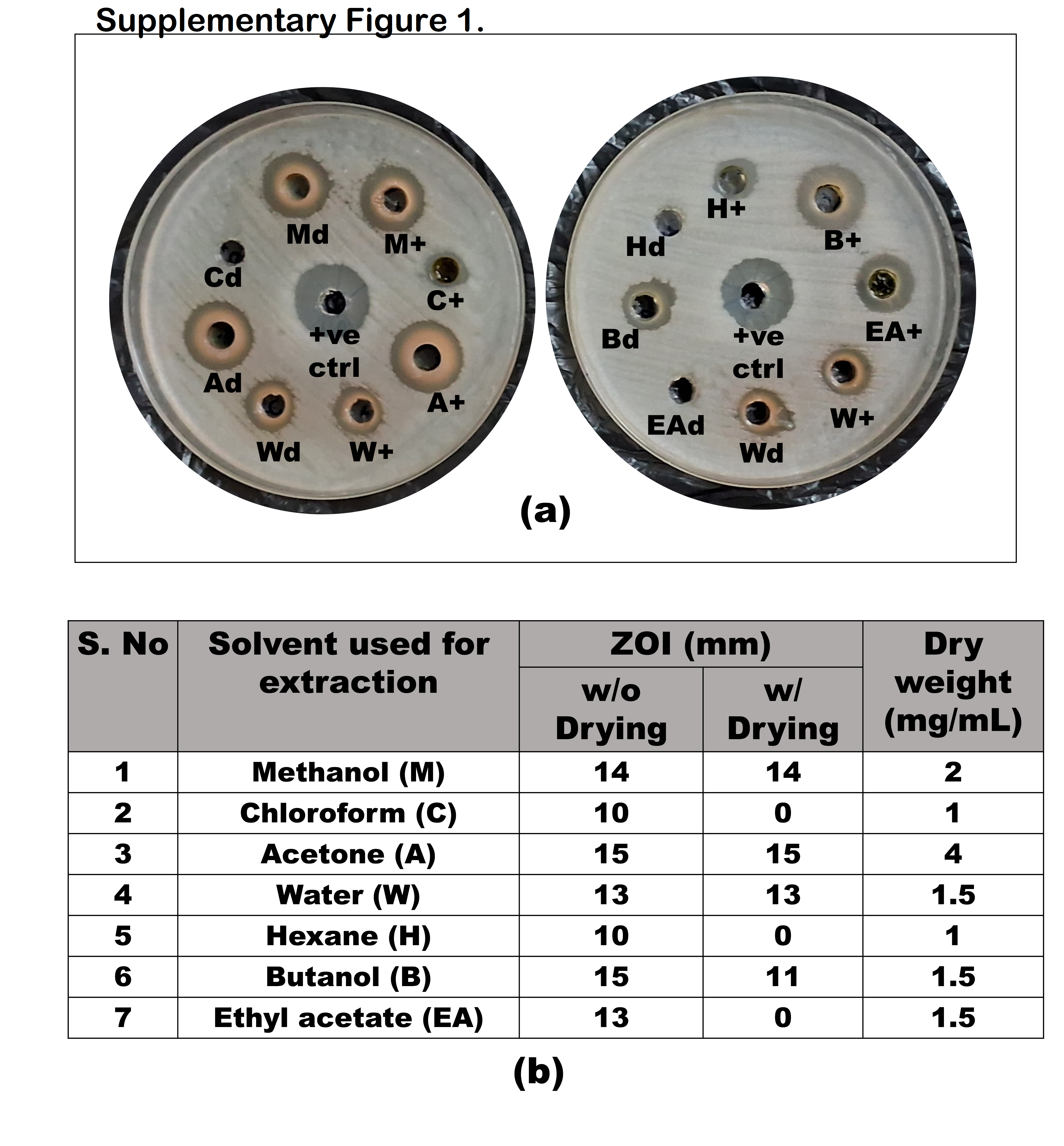

### Suppl. Figure 2

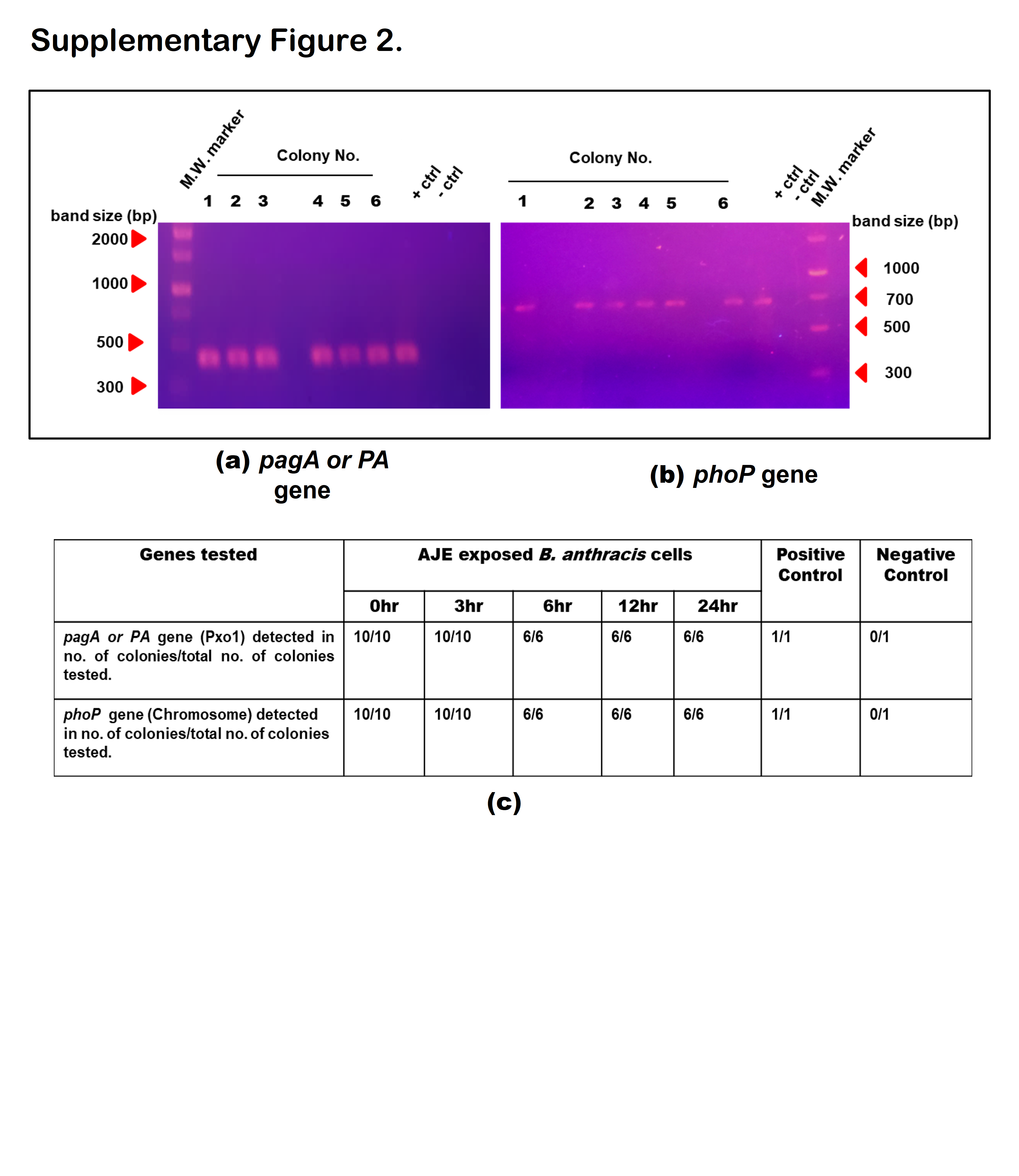

### Suppl. Figure 3

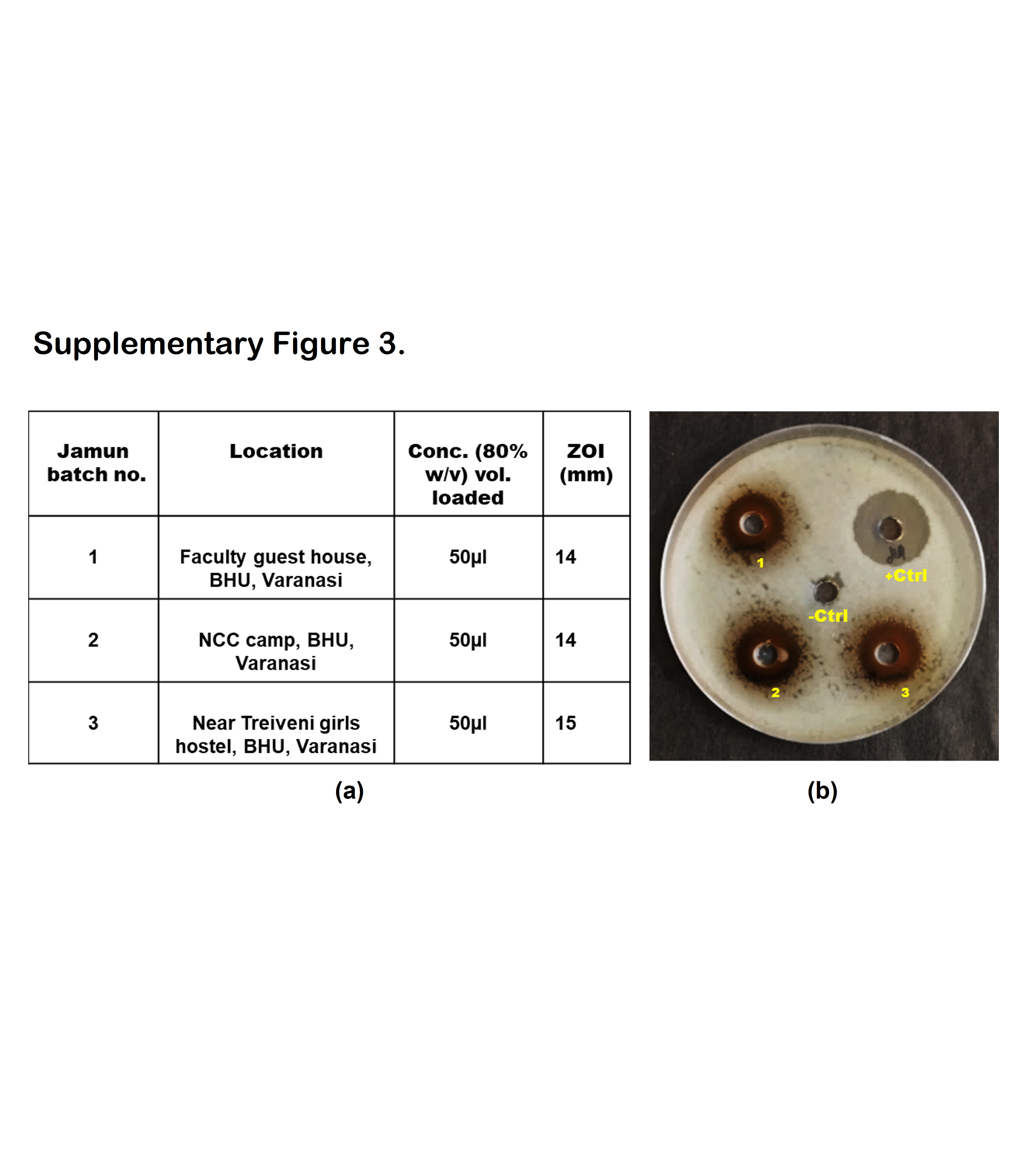

### Suppl. Figure 4

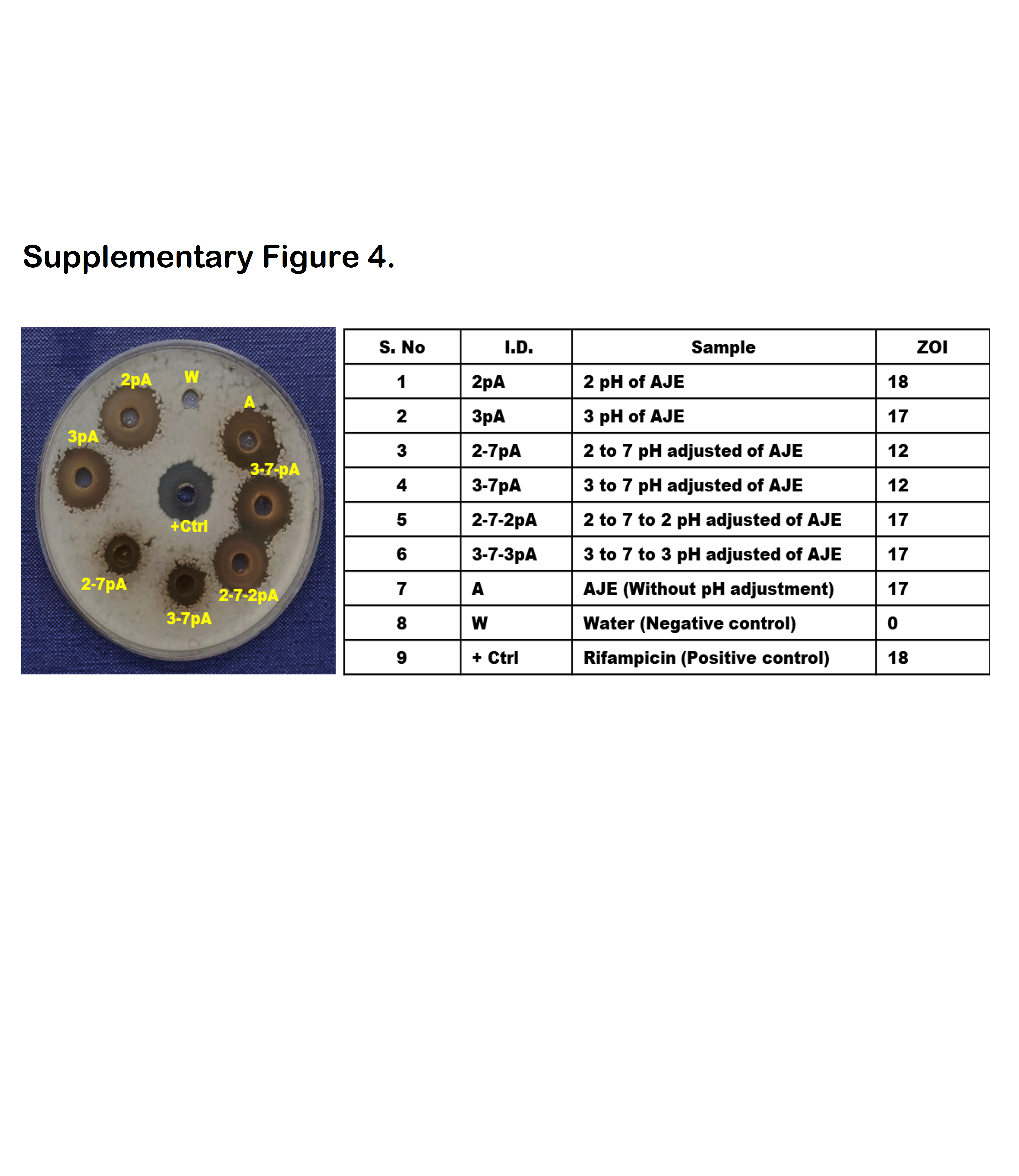

### Suppl. Figure 5

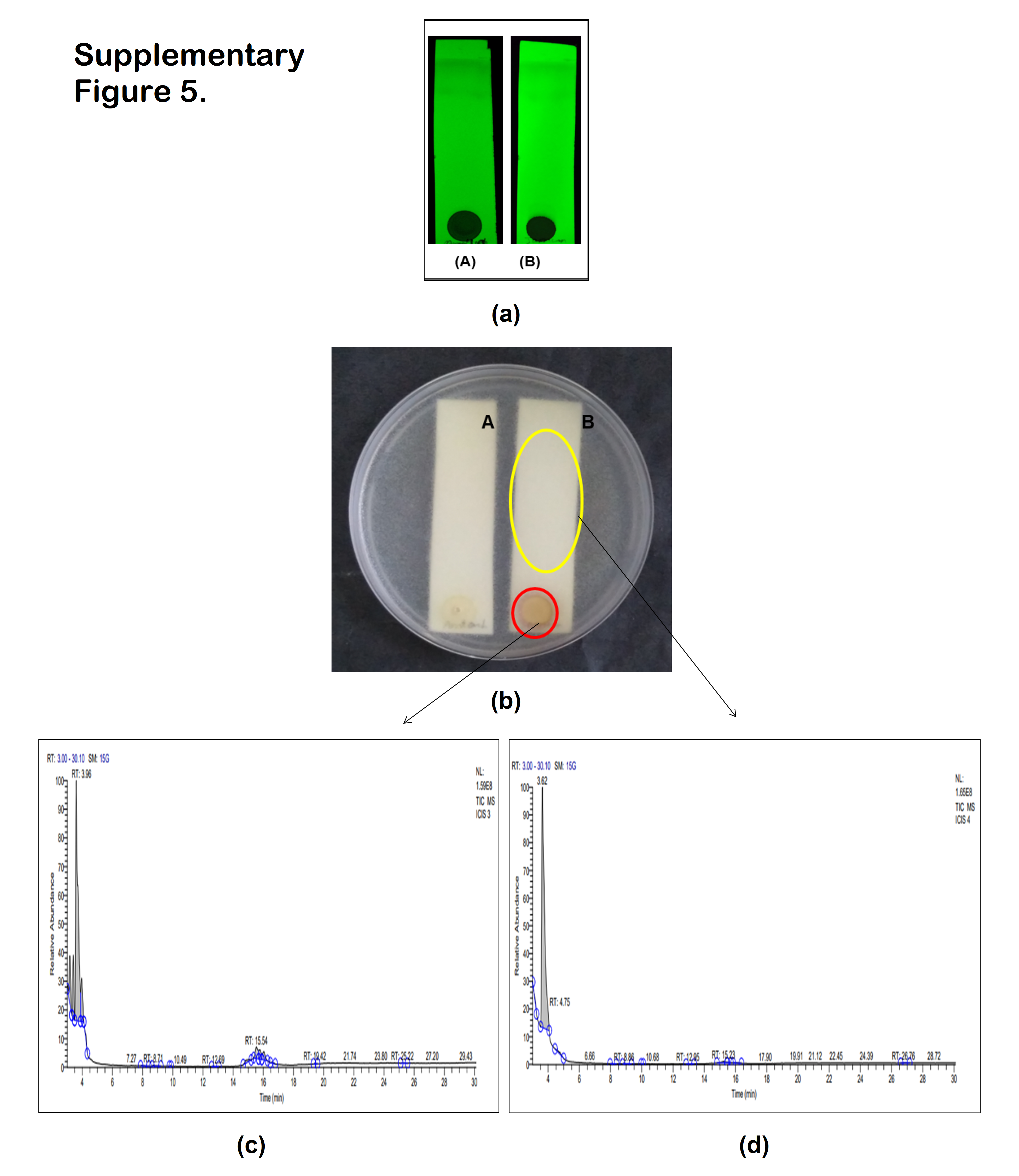

### Suppl. Figure 6

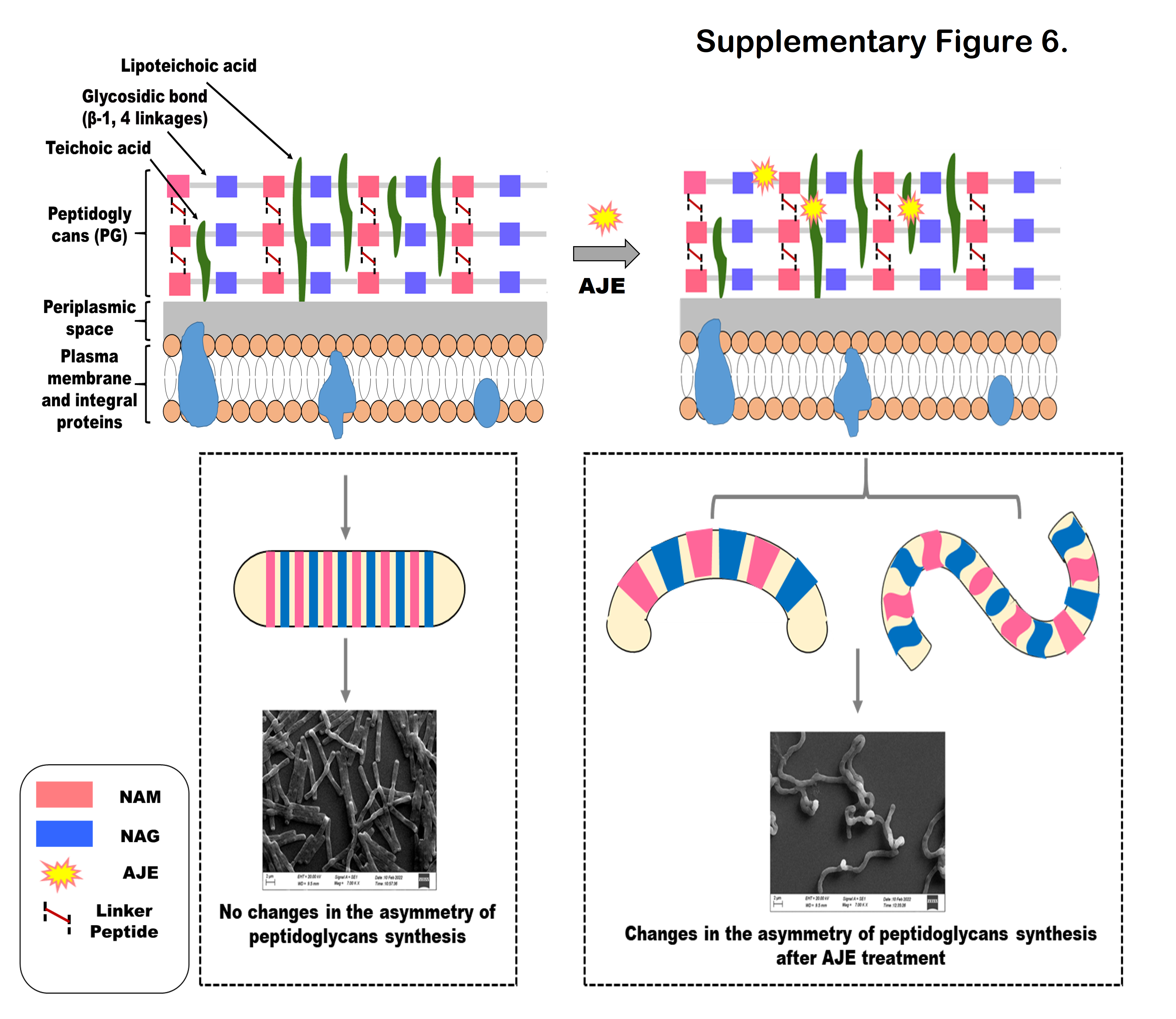
